# Stimulus presentation can enhance spiking irregularity across subcortical and cortical regions

**DOI:** 10.1101/2021.03.27.437321

**Authors:** Saleh Fayaz, Mohammad Amin Fakharian, Ali Ghazizadeh

**Affiliations:** Electrical Engineering Department, Sharif University of Technology, Tehran, Iran; School of Cognitive Sciences, Institute for Research in Fundamental Sciences, Tehran, Iran

**Keywords:** Neural variability, fano-factor, spiking irregularity, rate variability, network models, electrophysiology

## Abstract

Stimulus presentation is believed to quench neural response variability as measured by fano-factor (FF). However, the relative contribution of within trial spike irregularity (nΨ) and trial to trial rate variability (nRV) to FF reduction has remained elusive. Here, we introduce a principled approach for accurate estimation of variability components for a doubly stochastic point process which unlike previous methods allows for a time varying nΨ (aka ϕ). Notably, analysis across multiple subcortical and cortical areas showed across the board reduction in rate variability. However, unlike what was previously thought, spiking irregularity was not constant in time and was even enhanced in some regions abating the quench in the post-stimulus FF. Simulations confirmed plausibility of a time varying nΨ arising from within and between pool correlations of excitatory and inhibitory neural inputs. By accurate parsing of neural variability, our approach constrains candidate mechanisms that give rise to observed rate variability and spiking irregularity within brain regions.

## Introduction

While mean firing rate is widely used as a proxy of neural communication code, firing rate variance is also shown to play a significant role in neural coding and to serve as a diagnostic tool for distinguishing underlying neural mechanisms (A. K. Churchland et al., 2011; M. M. Churchland et al., 2006, 2010; M. M. Churchland & Abbott, 2012; Deco & Hugues, 2012; Goris et al., 2014; Shadlen & Newsome, 1998). In particular, neural response variability is believed to change during development and to be correlated with behavioral performance, stimuli conditions and to vary across different neural states (Lombardo et al., 2018; McIntosh et al., 2008; Mišić et al., 2010; Voytek et al., 2015; Waschke et al., 2021). Several studies have shown that spike count variability changes across different conditions in several cortical, subcortical, and cerebellar regions (A. K. Churchland et al., 2011; M. M. Churchland et al., 2010; Goris et al., 2014; Lombardo et al., 2018; Mitchell et al., 2007, 2009; Payne et al., 2019; Poland et al., 2019; Schölvinck et al., 2015). Importantly, numerous observations support a drop in neural response variability as measured by fano-factor (FF) in response to sensory or motor stimuli across cortical regions (M. M. Churchland et al., 2006, 2010; Rickert et al., 2009). The variation in neural spike count when viewed as a doubly stochastic process may be due to (1) the variation in generation of the spikes within the trials, (2) the variation between the trials of the same task (law of total variance). The first component is driven by within trial spiking irregularity (SI or ψ) and gives rise to the expected variance of the count (EVC) while the second component is driven by between trial rate variability (RV) and forms the variance of the expected count (VEC) (A. K. Churchland et al., 2011; Goris et al., 2014; Vinci et al., 2016). Despite this general understanding, a principled approach to estimate these two components of neural variability and to track their possibly concurrent changes in time is not developed yet. Consequently, it is not known how the widely reported post-stimulus reduction in FF in the brain arise from corresponding changes in RV and Ψ.

A previous attempt to estimate the RV component (A. K. Churchland et al., 2011) (VarCE method) showed that FF reduction is paralleled by the reduction in VEC (VarCE is an estimate of VEC). However, this approach assumed Ψ to be related to firing rate by a coefficient *ϕ* which was taken to be constant in time (see methods). Other more recent methods also followed the same assumption for spike count variability decomposition (Goris et al., 2014; Vinci et al., 2016). Clearly, such an assumption can result in gross misestimations of Ψ and subsequently VEC if there are temporal changes in *ϕ*. Even in the absence of such changes, VarCE estimates only the relative VEC up to arbitrary additive shifts. Moreover, the true nature and accurate estimation of ϕ also remains unaddressed.

Here, by invoking minimal assumptions from doubly stochastic renewal theory, we introduce an approach that explicitly relates FF to normalized RV and Ψ (nRV and nΨ) and reveals the nature of both components in relation to the first and second moments of inter-spike-interval distribution (ISI) (FF asymptote or FFA method). In addition, a second approach, adapted and re-purposed from a previously introduced method for estimating components of cross neural spike count correlation (Vinci et al., 2016) is also used. Results from both methods show that unlike what was previously assumed, nΨ robustly changed along with nRV during the stimulus presentation across several subcortical and cortical regions. While the post-stimulus nRV showed a ubiquitous change across all areas examined, post-stimulus nΨ change was more heterogenous: decreasing in some (mostly cortical) areas and increasing in others (mostly subcortical). Moreover, while we found the post-stimulus reduction in FF to be mostly due to nRV decrease, the concurrent change in nΨ was shown to convey critical information regarding the network state. Our results warn against the constant nΨ (*ϕ*) assumption when studying post-stimulus response variability and cautions against strong claims to ubiquitous response variability quench across the brain. The heterogeneity in patterns of nRV and nΨ during an event constrains underlying neural mechanisms differently for different regions.

## Results

### Parsing neural response variability components: the method

A stochastic process with deterministic parameters would induce variability in the spike count, i.e. point-process variability. Moreover, stochastic selection of such parameters between different trials would impose variability across trials (doubly stochastic process). We assume a doubly stochastic renewal process in which inter-spike interval (ISI) parameters change probabilistically across trials. In this case, sources of spike count variability can be decomposed into at least two components using the law of total variance in the following form:

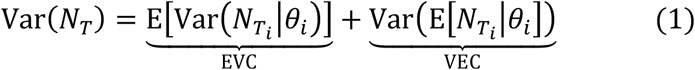

Where *N_T_* denotes the spike count in the time-bin (T) and *θ_i_* represent ISI parameters in trial i. Here the first term denotes the expected variance of spike count (EVC) due to spiking irregularity (Ψ) within a trial given constant parameters, and the second term denotes variance of expected count (VEC) due to rate variability (RV). In the most general sense accurate estimation of these two components is not feasible as we only can measure one sample of N_T_i__; hence, no estimates of the latent variable *θ_i_* is explicitly known. However, progress can be made for renewal processes which has been widely used as models of spiking neurons (Fisch et al., 2012; Koyama & Kostal, 2014; Lansky et al., 2016; Nawrot et al., 2008; Omi & Shinomoto, 2011; Ostojic, 2011; Pipa et al., 2013; Rajdl et al., 2020; Shimokawa et al., 2010; Shinomoto et al., 2005). Fano-Factor estimation shows a functionality of time-bin (T); In particular it can be shown that the fano-factor (FF) of a renewal doubly stochastic process in sufficiently large time-bin follows the formula below (see methods for details):

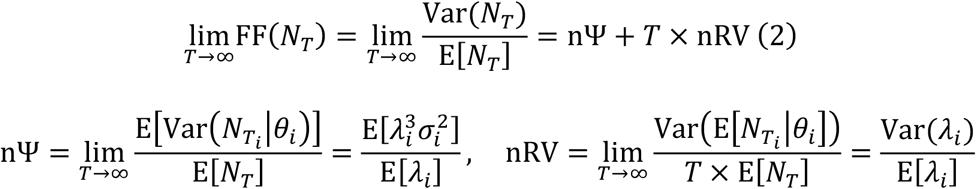

Where 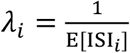 and 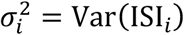 and ISI_*i*_ is the random variable representing the inter-spike intervals drawn from a given distribution whose parameters can vary across trials. nΨ and nRV represent normalized within trial spiking irregularity (Ψ) and between trials rate variability (RV) respectively (see methods). As can be seen in equation 2, nΨ and nRV depend solely on the first and second moments of ISI distribution for renewal processes. Note that since FF in equation 2 is a linear sum of these two components the changes in FF may reflect change in either or both components.

According to equation 2, FF is asymptotically a linear function of time-bin. It includes an intercept which represents normalized within trial spiking irregularity (nΨ aka *ϕ*) and a slope which represents normalized between trials rate variability (nRV). Therefore, a linear regression of FF against time in the asymptotic part would provide both components of FF. We call this method, the FFA method. For a simple stochastic point process with no trial-to-trial variation in ISI parameters, FF should be constant for sufficiently large time-bins. Figure 1a shows that indeed after a short transient, FF converges to the theoretical prediction based on the parameters of the ISI distribution (i.e. it is 1 for Poisson point process and is 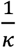 for Gamma(*κ,θ*), see methods). Consistent with the well-known equality of FF and CV^2^ in stationary renewal process (Buckland & Cox, 1964), it can be shown that - even for non-stationary processes - nΨ can be approximated by 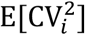 where CV_*i*_ is the coefficient of variation in trial *i* (see methods). This is intuitively encouraging since local CV measures have been widely used as a measure of spiking irregularity (Miura et al., 2007; Nawrot et al., 2008; Payne et al., 2019; Ponce-Alvarez et al., 2010). On the other hand, for a doubly stochastic process with trial-to-trial variation in rate, FF increases in an asymptotically linear fashion as a function of time-bin. Note that in this case the intercept of the line which represents nΨ remains unchanged as long as the average spiking irregularity (e.g. *κ* in Gamma(*κ, θ*)) remains unchanged across trials (Figure 1a). Notably, this result holds for any renewal process regardless of the shape of ISI distribution as illustrated for exponential, gamma and inverse gaussian probability density functions (PDFs). Also note that while in theory FF asymptotic behavior is at infinity limits of time-bin, in practice the linear asymptotic behavior may be observable with much smaller time-bins depending on the firing rate of the neuron (Figure 1a, S1).

**Figure 1.**
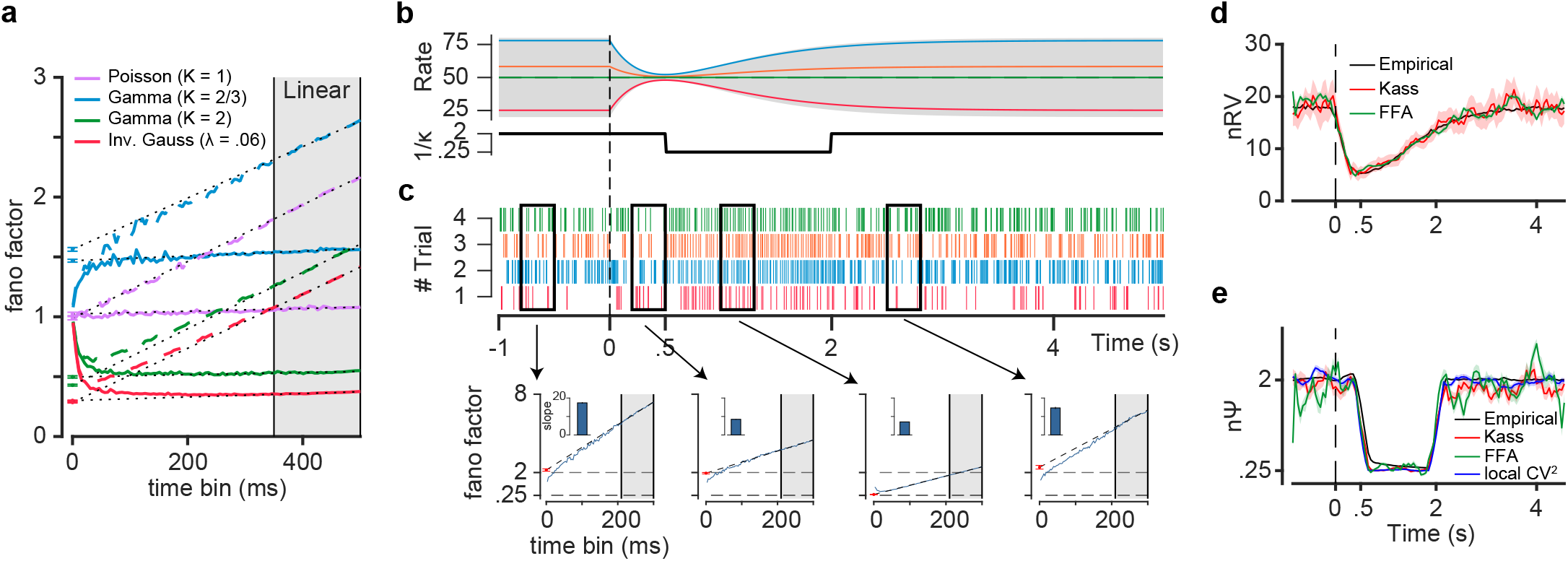
Parsing components of variability in doubly stochastic renewal processes in time. (a) Fano factor (FF) as function of time bin ([0,1] ratios of time-bin) for simple (solid lines) and doubly stochastic (dashed lines) renewal processes with different ISI distributions (exponential, gamma and inverse gaussian). The asymptotic part of FF for large time bins (gray shaded area) is used for independent estimates of normalized spiking irregularity (nΨ) as the intercept and the normalized rate variability (nRV) as the slope. Note the non-zero identical slopes only for doubly stochastic processes (*λ_i_* ~ *N*(*μ* = 50Hz, *σ* = 10Hz)). (b) Rate parameter 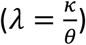 and inverse of shape parameter 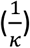 for a doubly stochastic gamma process as a function of time. Rate parameter was variable across trials according to 50 + *n_i_* × (1 – 5*t*_+_*e*^−2*t*_+_^) where *n_i_~N*(0,30) and *t*_+_ is time for positive values and zero otherwise. (c) top, raster plot realization of color-coded trials from the stochastic gamma process (rate of each trial is indicated in panel b), bottom, FF as a function of time bin within the specified time-bin (300ms), blue curve represents FF estimates as a function of time. Dashed line shows extrapolation of fitted line in linear part; red dot represents intercept. Slopes of each line are indicated by bar plots. (d) nRV estimates made empirically by assuming access to a large number of realizations given constant parameter (black) along with estimates using smoothed FFA (green) and modified Kass (red) methods. (e) same format as d but for nΨ estimates. Blue trace represents 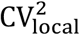 estimates. All error bars indicating the sem.

Using the FFA method, one would be able to track the temporal fluctuations in nΨ and nRV during a trial using a sliding time window. Figure 1b shows a simulated doubly stochastic spike process with concurrent variations in Ψ and RV across time. Each trial was generated from a renewal point process with ISIs following a two-parameter Gamma(*κ,θ*) distribution. The process had two sources of variability: Firstly, the rate in each trial was 50 + *n_i_* × (1 – 5*t*_+_*e*^−2*t*^+) where *n_i_*~*N*(*μ* = 0, *σ* = 30) and *t*_+_ is time for positive values and zero otherwise. This led to a constant mean firing of 50Hz across trials with a variance that changed during the trial (Fig. 1b top). This variation of rate around the 50Hz mean is responsible for RV. Secondly, in each trial, spikes were generated with the variability inherent to a gamma point process. In this case the parameter *κ* is inversely proportional to spiking irregularity and was set to change in time (e.g. changes in 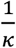 from 2 to 0.25 to 2 in time). Figure 1c shows spiking realization of this doubly stochastic gamma process across four trials each corresponding to a given rate dynamic. Interestingly, FF intercept and slope in four selected 300ms epochs closely paralleled the expected changes in nΨ and nRV during the trial. Indeed, the slope showed a reduction concurrent with the reduction of rate variability after time zero while the intercept independently followed changes in 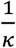 that was delayed (onset asynchrony) compared to RV (going from 2 to 0.25 and back to 2).

The FFA method can be used to track the temporal dynamics of nΨ and nRV for this simulated neuron (Figure 1d-e). As can be seen, the FFA method uncovers concurrent reduction in both spiking irregularity (nΨ) and rate variability (nRV) as well as the asynchrony in this reduction (nΨ reduction happening after nRV). The overlaid empirical estimate shows the best possible estimate from the data if one had access to a large number of repetitions for each trial with the same ISI parameters (say>200 realizations of the point process with the same *θ_i_*).

While the FFA method has nice theoretical underpinnings and pedagogical value, in practice fitting the asymptotic behavior of FF as a function of time-bin can be noisy especially for the intercept (small change in slope translating to large swings in the intercept). Therefore, we used a more robust estimator of nΨ and nRV by adapting a previously introduced method (Vinci et al., 2016) and allowing for variations of nΨ in time (see methods for details). This modified method which we refer to as the Kass method enables us to estimate concurrent changes in nΨ and nRV with higher fidelity compared to FFA (Figure 1d-e). Thus, along with the FFA slope and intercepts for each region, in practice the Kass method will be used to track sources of variability in time throughout this study. Note that both FFA and Kass methods estimation require sufficiently large number of trials and spikes per time-bin to give accurate estimates of nΨ and nRV. However as noted, the Kass estimate seems to be more robust compared to FFA across a range of parameters (Figure S1).

Normalized point-process variability (nPPV) which represents the inherent randomness of spike generation can be quantified using a measure which we refer to as 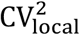 that estimates the inverse of shape parameter *κ*. This measure which is based on a local CV measure (known as CV2) is shown to provide robust and unbiased estimation of shape parameter (Holt et al., 1996; Pachitariu et al., 2010) (see methods). In the current simulated example, there are no other sources of nΨ other than nPPV. Hence, as illustrated in figure 1e, 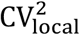 also matches the nΨ estimate both tracking well the changes in 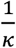. In the following sections, we will show that in general nPPV is the lower bound of nΨ and that in addition, spiking irregularity is increased by other non-stationarities during the trial.

The fact that the firing rate of real neurons can change during the relatively large time-bins can raise a concern about the inflated estimates in nΨ. However, our analysis and simulation results show that as long as the spiking irregularity remains unchanged during this time period our estimates of nΨ is largely robust to the underlying rate changes (Figure S2).

### Post-stimulus enhancement of nΨ and reduction of nRV in cerebellum

Cerebellar neurons show sizeable changes in their variability in response to different conditions (Häusser & Clark, 1997; Payne et al., 2019; Shin et al., 2007; ten Brinke, Heiney, Wang, Proietti-Onori, Boele, Bakermans, Medina, Gao, & De Zeeuw, 2017). We used our method to parse rate variability and spiking irregularity across neurons (n=268) recorded from Interposed Nucleus (IpN) of mice during the Pavlovian eyeblink conditioning task (ten Brinke, Heiney, Wang, Proietti-Onori, Boele, Bakermans, Medina, Gao, & De Zeeuw, 2017). Figure 2a shows peristimulus time histogram (PSTH) response of an example IpN neuron time locked to the conditioned stimulus (CS) onset (LED light). 250ms after the CS onset the unconditioned stimulus (US: air-puff) was delivered. As can be seen the example neuron showed strong excitation following CS onset and just prior to the US delivery. Figure 2b top shows an initial decrease in the average FF followed by a sustained increase. As can be seen, this temporal pattern was largely due to very similar pattern of change in rate variability (nRV). This pattern of change in nRV and in particular its rise after about 500ms from CS can be easily verified by looking at the spiking raster plot (Figure 2a bottom). Interestingly, despite the initial decrease in nRV and FF, nΨ showed a robust post stimulus increase. Notably the increase in nΨ was only partially due to the increase in nPPV as measured by 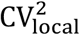 pointing to extra factors elevating the observed spiking irregularity.

**Figure 2.**
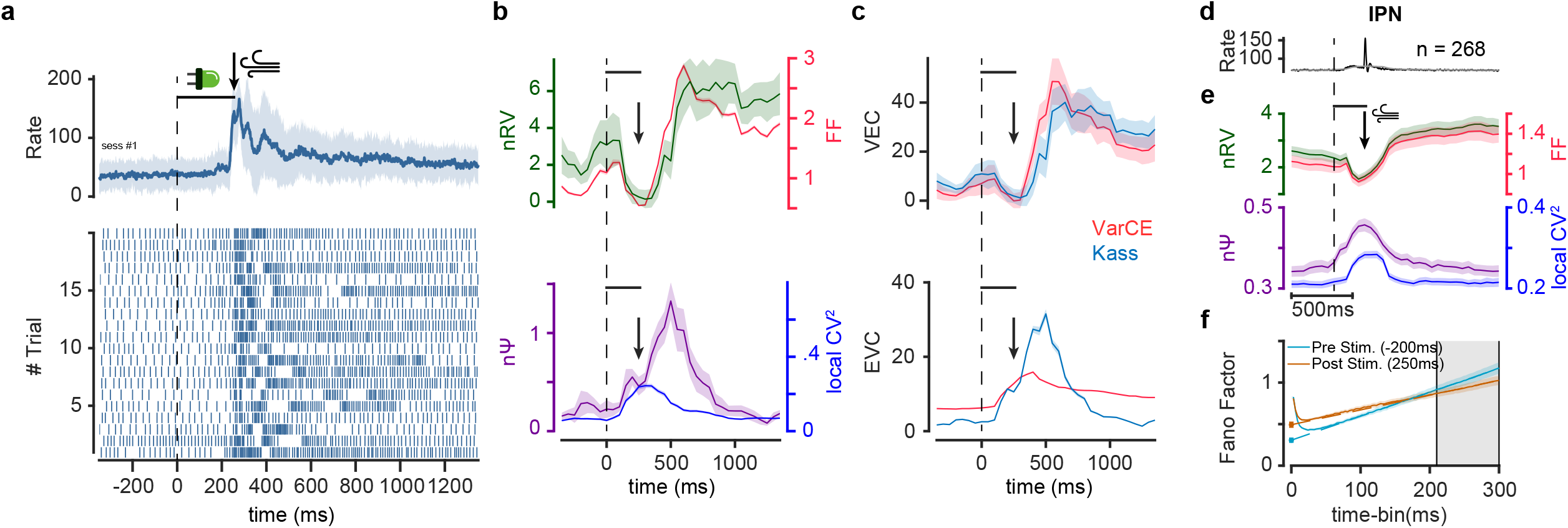
Post-stimulus changes in rate variability and enhancement of spiking irregularity in cerebellum. (a) Example IpN neuron PSTH with standard deviation as the shaded area (top) and raster plot of 20 example trials (bottom) aligned to the conditioned stimulus (CS) onset (LED light) followed by unconditioned stimulus (US, air-puff) in 250ms. (b) Post-stimulus FF and nRV showing an initial decrease followed by sustained increase (top) and post-stimulus nΨ showing a robust increase. 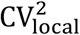 also shows a moderate increase following stimulus onset. (f) VEC and EVC estimations using the Kass vs VarCE method (top and bottom, respectively). VarCE overestimates the rise in VEC (related to rate variability) and underestimates EVC (related to spiking irregularity). (d) Average PSTH of IpN neural population (n=268) neurons. Black and gray traces show PSTHs with 10ms and 300ms bin-widths, respectively. Longer bin-width is equal to time-bin used for variability estimation in the Kass and FFA methods. (e) Average FF and nRV (top) and nΨ and 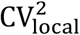 (bottom) as a function of time within the sliding time-bin (300ms, 50ms sliding). Dashed line indicates CS onset, horizontal line indicates CS on period and arrow indicates US onset. (f) FF as a function of time bin within time-bin (300ms) centered on pre- and post-stimulus onset periods (−200 and 250ms respectively). Gray area indicates the linear zone at which asymptotic FF is estimated. The increase in intercept and the decrease in slope is consistent with changes in nΨ and nRV respectively. Error bars are sem.

Given the observed changes in nΨ, methods that assume a constant *ϕ* (i.e. constant nΨ or spiking irregularity) can misestimate nRV. For instance, the VarCE method which attempts to estimate VEC by finding the largest constant *ϕ* (nΨ) that keeps VEC positive during the trial, can become error prone:

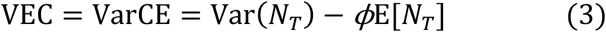

The VarCE method is also based on parsing sources of variability using the law of total variance in which VarCE and *ϕ* here are measures of VEC and nΨ, respectively. Note that even in cases when *ϕ* is constant, VarCE can only measure VEC up to additive shifts (Figure S3a). More importantly, if the true *ϕ* changes and in the absence of changes in VEC, VarCE erroneously reports changes in VEC (Figure S3b). In the case of our cerebellar neural example, VarCE estimated the trends of change in VEC accurately but overestimated it particularly during the time when nΨ was changing (Figure 2c top). More importantly, since estimated *ϕ* in VarCE was constant, it grossly misestimated the component of variability that was due to spiking irregularity VEC (i.e *ϕ*E[*N_T_*]) (Figure 2c bottom).

Similar to the neural example shown, the pattern of FF reduction and subsequent increase was also observable across the cerebellar IpN population and seemed to be driven primarily by the temporal dynamics in nRV. Once again during the same period nΨ showed a robust post stimulus increase which was accompanied by concurrent increase in nPPV which was nevertheless only partially explained the increase observed in nΨ. The post stimulus reduction in nRV and increase in nΨ can also be verified using the FFA method by looking at the linear asymptotic behavior of FF which showed intercept increase and slope decreased after the stimulus onset.

Simulation results show that in the absence of within trial non-stationarities, nΨ will be the same as nPPV as measured by 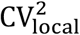 regardless of particular ISI distributions used (Figure 3a-b) and the existence of rate variability (Figure 3c). Both FFA and Kass methods track the ideal values of nΨ and nRV with good accuracies in such cases (Figure 3a-c). So, what can cause the mismatch between nΨ and nPPV seen in Figure 2b, e? Figure 3d-f shows three possible scenarios in which nΨ can be inflated beyond nPPV including bursting (Ponce-Alvarez et al., 2010), rate noise (Ermentrout et al., 2008) or rate switching (Deco & Hugues, 2012; Song et al., 2018) within the time-bin. All of these effects create stochastic non-stationarities during the time-bin and can increase spiking irregularity. In the case of rate switching with low transition probability there is also a concurrent increase in between trial rate variability which is well-tracked by both FFA and Kass methods. Note that in all cases 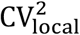 proves to be a good estimator of nPPV reflecting its robustness to rate fluctuations within and between the course of the trial. Thus, one may use 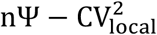 as a measure of spike irregularity induced by sources other than the point process itself which may also interact with nPPV to increase nΨ.

**Figure 3.**
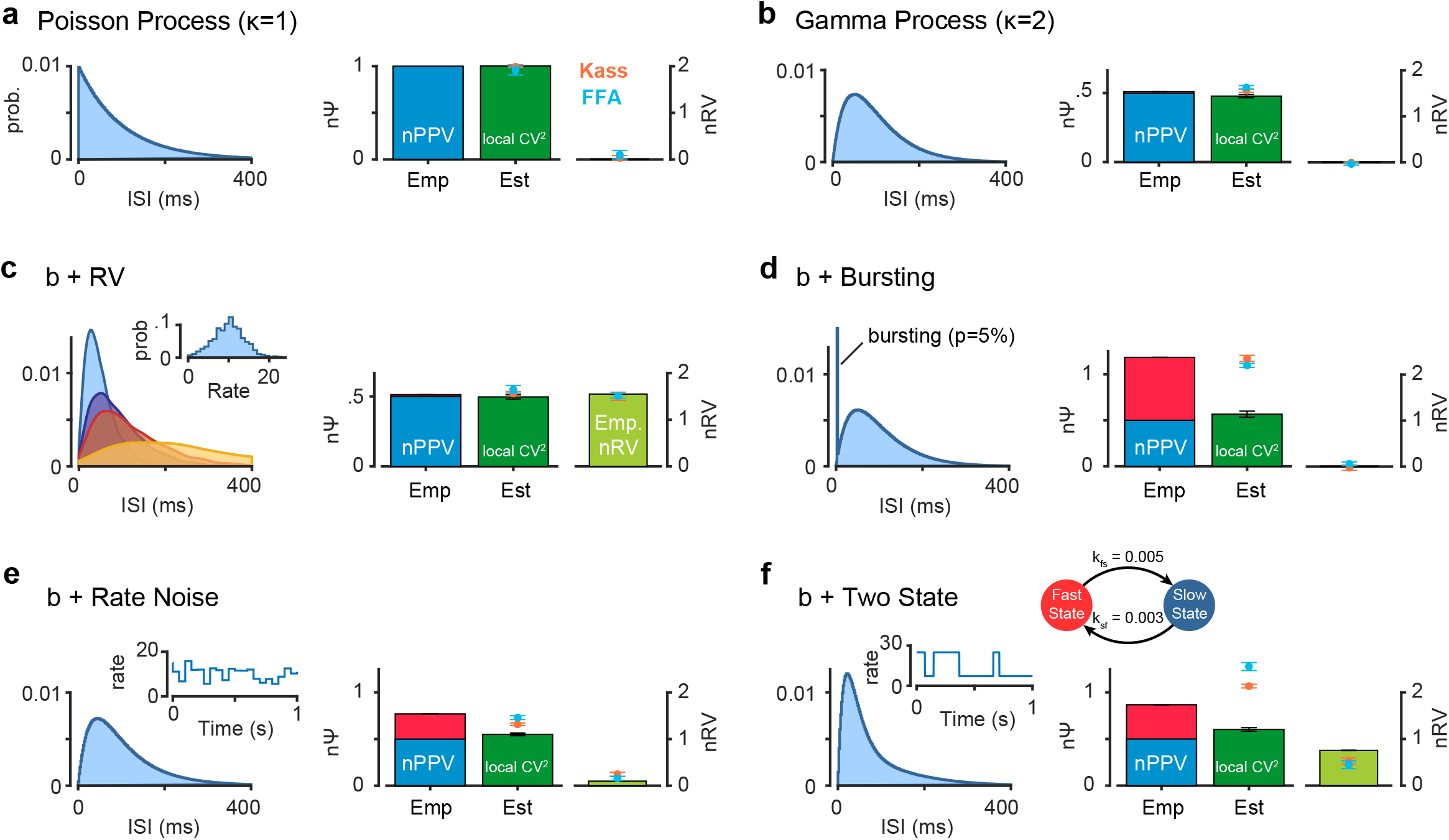
Within trial non-stationarities as well as normalized point process variability (nPPV) contribute to spiking irregularity (nΨ). Empirical estimations of nΨ and nRV along with estimations made by FFA, Kass methods. Theoretical nPPV and its estimate using 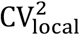 is also shown. (a) Poisson process with rate 10Hz with no rate variability. (b) Gamma process (*κ* = 2) with no rate variability. (c) same as b but doubly stochastic with rate variability. (d) same as b but with bursting imposed with probability 0.05 on a given spike (5 spikes with 3ms ISI per burst). (e) same as b but with within trial white noise added to the constant rate (~ *N*(0,3) and Δ*t* = 50ms, see methods). (f) same as b but with within trial state switching between high and low-rates imposed (see methods). Manipulations shown in d-f can increase nΨ beyond irregularity imposed by nPPV (red portion in empirical). In all cases 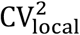 is high fidelity estimated of nPPV. State switching can also result in increased nRV. FFA and Kass methods provide good estimate of nΨ and nRV in all scenerios.

### Post-stimulus enhancement of nΨ canceling reduction of nRV in substantia nigra

Next, we checked response variability in another subcortical area namely substantia nigra reticula (SNr). This region is particularly informative for our analysis since here individual neurons can show large dynamic range of excitatory or inhibitory responses to visual objects based on their past value memory (excitation to low value objects and inhibition to high value objects). Figure 4a-b show responses of example neurons to high value and low value objects, respectively. For the neuron with inhibitory response a post stimulus reduction in FF was observed as expected. Similar to what was observed in cerebellum this decrease was largely due to a drop in post-stimulus rate variability (nRV) rather than spiking irregularity (nΨ) which instead showed a robust post stimulus increase (Figure 4c). Interestingly, the neuron with excitatory response also showed a drop in nRV. However, this drop was more than offset by the increase in the spiking irregularity such that post stimulus FF in this neuron did not show the usually reported quench in spiking variability (Figure 4d). As expected, the radical changes in nΨ in both neurons meant that the estimates of VEC using VarCE were erroneous (Figure 4e-f). Moreover, there was a gross misestimation of EVC using VarCE especially in the case of the neuron with the inhibitory response. This is because methods with constant *ϕ* assumption (equivalently constant nΨ), inevitably predict a drop in Ψ for inhibitory responses which turned to be the opposite of what was observed for this neuron (Figure 4e).

**Figure 4.**
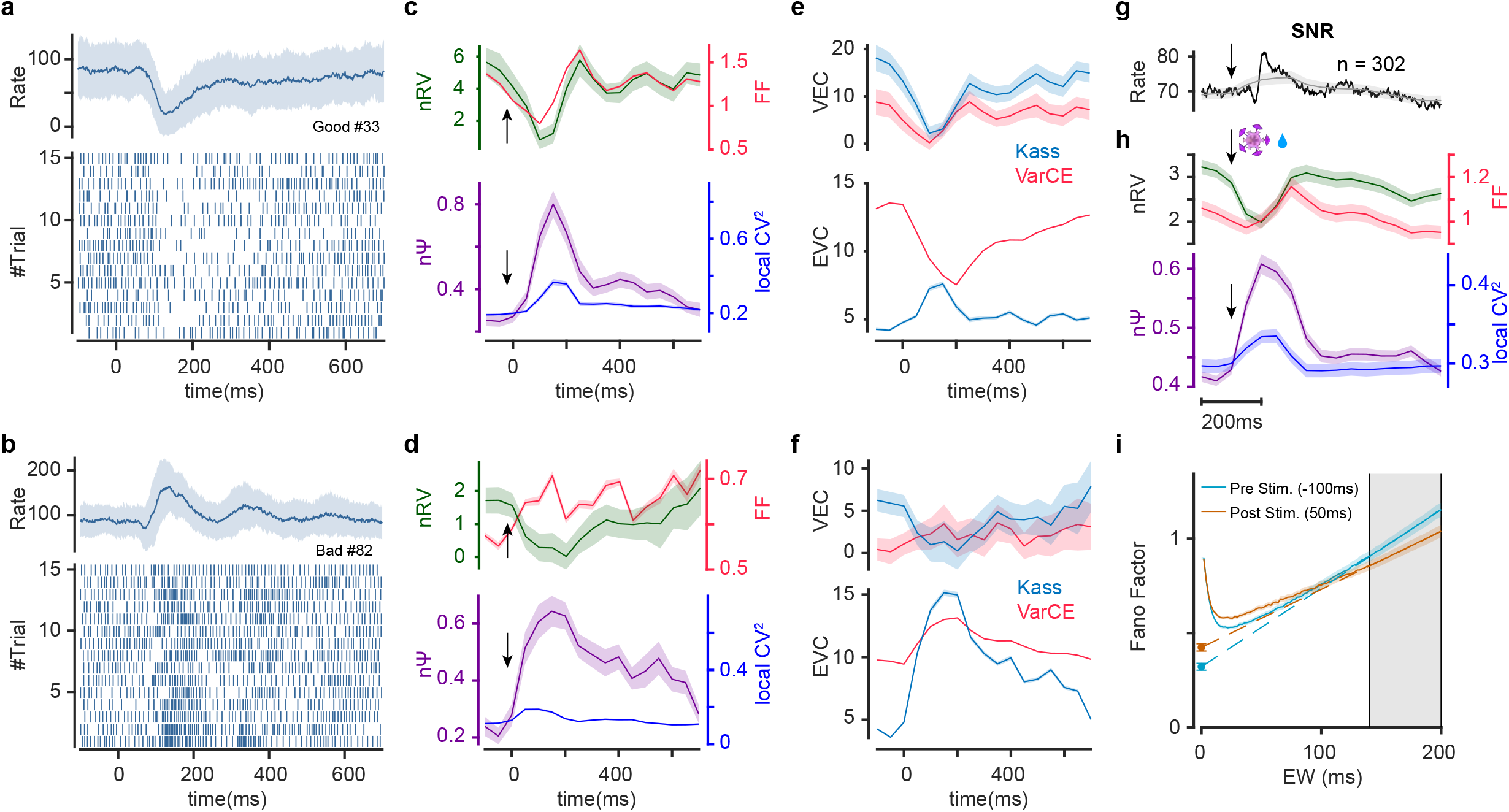
Post-stimulus drop in rate variability is compensated by concurrent rise of spiking irregularity in SNr. (a-f) Same format as Figure 2a-c but for one example SNr neuron showing inhibition (a, c, e) and another example SNr neuron showing excitation (b, d, f) to object onset. Raster plots show 15 sample trials for each neuron. (g-i) Same format as Figure 2d-f showing population average of 302 SNr neurons.

Across the population of recorded neurons, stimulus onset was associated with a drop in nRV, on the other hand, nΨ showed a robust increase in response to the stimulus onset revealing enhanced spiking irregularity (Figure 4g-h). Similar to the cerebellar neuron, population 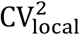 also increased following the stimulus onset suggesting a concurrent increase in nPPV which nevertheless failed short of explaining the larger increase in nΨ (Figure 4h). The decrease in nRV and increase in nΨ can be visualized by the FFA method as well (Figure 4i). Note that unlike the cerebellar IpN neurons due to relatively large size of nΨ, dynamics of the FF and is dissimilar for this region (Figure S4). Indeed, SNr is one brain region which also violates the well accepted reduction of post stimulus FF (M. M. Churchland et al., 2010).

As shown in equation 2, these results were obtained using normalized measures of between and within trial variabilities to control for firing rate change during the stimulus. Nevertheless, and to ensure the robustness of our findings to firing rate changes during the trial (a confounder), we validated our findings using two other approaches: (1) by using the mean-matching method which chooses subsets of neurons at each time point with the constraint of similar rate distributions (M. M. Churchland et al., 2010) and (2) by repeating the analysis separately for neurons in different response types of excitatory, inhibitory and null post-stimulus responses (see methods). Both scenarios resulted in essentially the same behavior for nΨ and nRV and relatively stable FF during the trial for the SNr population (figure S5).

### Stimulus evoked changes in nΨ and nRV across cortical areas

It was previously shown that stimulus onset tends to reduce variability in cortical neural responses (M. M. Churchland et al., 2010). However, as stated previously due to lack of reliable methods for parsing the sources of spiking variability, it is not known how nΨ and nRV are affected by the stimulus onset. Here, we applied our methods on multiple cortical regions including areas V1, V2, middle-temporal (MT), premotor cortex (PMd) and ventrolateral prefrontal cortex (vlPFC) recorded in macaque monkeys by different groups (M. M. Churchland et al., 2010; Ghazizadeh et al., 2018; Zandvakili & Kohn, 2015). As can be seen stimulus presentation or motor action caused a sizeable increase in average firing rate and a decrease in FF of all examined areas (Figure 5a). Moreover, in all cases the reduction in FF was concurrent with stimulus related reduction of nRV (Figure 5b). However almost all areas showed concurrent changes in nΨ as well. With the exception of vlPFC, the spiking irregularity as measured by nΨ decreased during stimulus presentation across cortical regions. In vlPFC a transient post-stimulus increase in nΨ was observed. Note that despite the transient increase in nΨ in vlPFC, its FF was still quenched during stimulus presentation due to larger size of nRV compared to nΨ (note the y-scales, Figure S4). Indeed, in all areas one can verify that nΨ and nRV scaled by time-bin sum up to explain the total FF fluctuations across time (Figure S4). The changes in nΨ and nRV during stimulus on and off is also evident by looking at the intercept and slope of the FF in the time-bin (Figure 5c). Note that in regions with lower overall firing rate, larger time-bin is required for the asymptotic behavior of FF to emerge (compare 1000ms time-bin in V1 with 200ms time-bin in vlPFC). It is thus possible that for regions with low firing rate, fast temporal dynamics in nΨ and nRV not be detectable with low number of trials and large time-bins (such as transient increases similar to vlPFC). The relationship between nPPV and nΨ was more heterogenous between regions. While nPPV hardly changed in V1 and V2, it decreased during stimulus presentation in MT and vlPFC. Once again nPPV changes were relatively minimal and most of the changes in spiking irregularity was not due to nPPV (Figure 5b).

**Figure 5.**
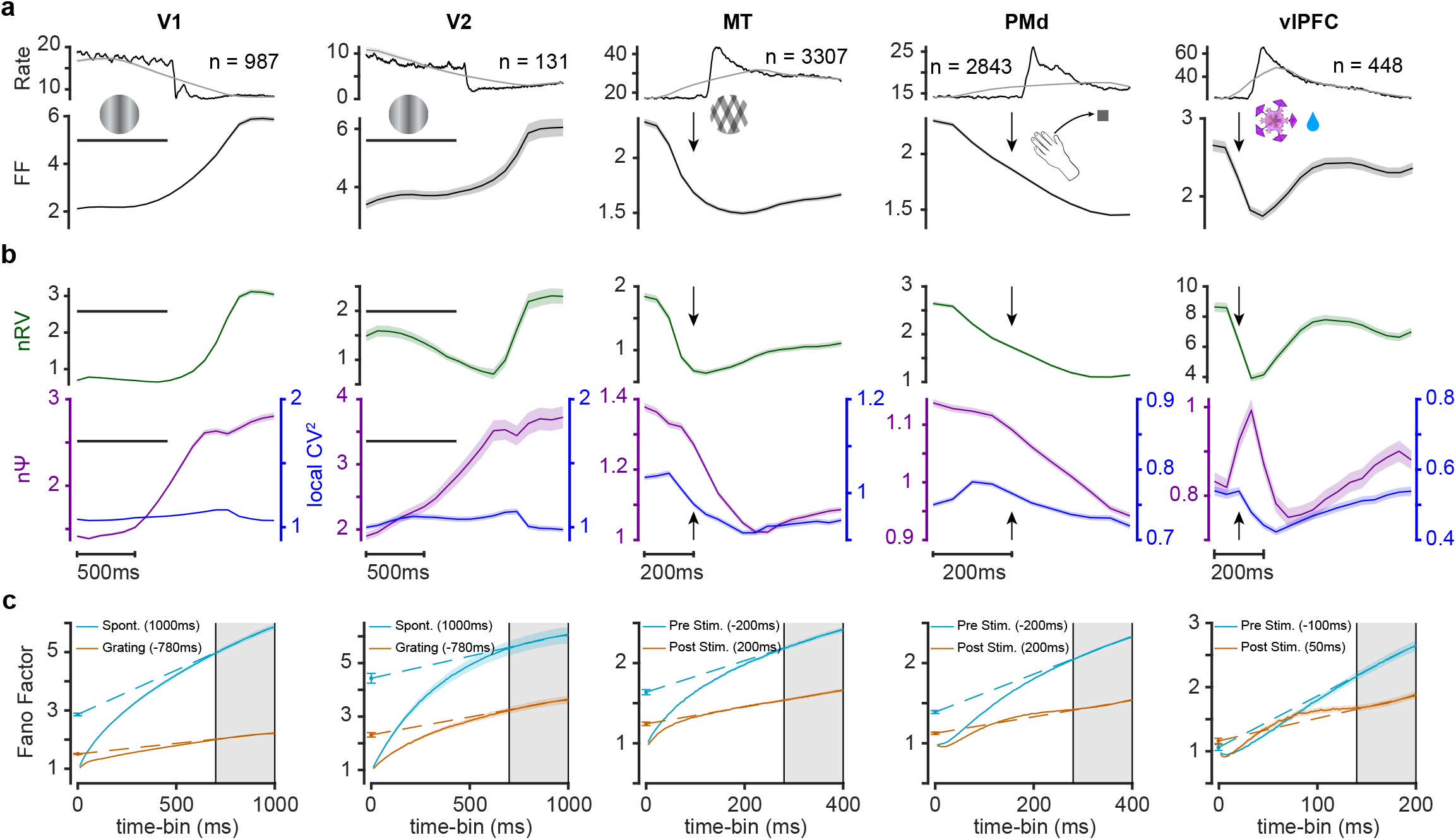
Concurrent changes in rate variability and spiking irregularity during stimulus presentation or movement execution across the cortex. (a) Average population PSTH across 5 different cortical regions responding to the stimulus presentation (areas V1, V2, MT, vlPFC) or movement execution (PMd). Black and gray traces show PSTHs with 10ms and EW bin-widths, respectively (time-bin = 1000,1000,400,400,200ms sliding by 100ms for V1/2 and 50ms for the rest respectively). Longer bin-width is equal to time-bin used for variability estimation in the Kass and FFA method (top), and corresponding change of FF during the trial (bottom). Horizontal line indicates stimulus on period and arrow indicates stimulus onsets. (b) Concurrent temporal changes in nRV (top) and in nΨ (bottom) across cortical regions. Temporal dynamics of 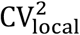 is plotted alongside nΨ. (c) Same format as Figure 2f for each cortical area pre- and post-sensory or movement event.

These results were further corroborated using mean matching across the full population as well as separately within clusters of neural responses showing excitatory, inhibitory or null response across all the cortical regions (Figure S6 mean-matching and Figure S7 response clustering). In particular the transient increase in nΨ in vlPFC was observed in all cases.

### Different schemes for coordinated activity in the presynaptic network induce independent changes in nΨ and nRV

As mentioned previously, accurate estimation of components of neural variability constrains the viable neural mechanisms that underlie the observed neural responses and can serve as a valuable tool for cross region comparisons (A. K. Churchland et al., 2011; Deco & Hugues, 2012; Gómez-Laberge et al., 2016; Rajan et al., 2010; Shadlen & Newsome, 1994; Softky & Koch, 1993). CV of ISIs was previously reported to change as a function of correlations in pre-synaptic inputs both theoretically and using simulations (Salinas & Sejnowski, 2000). However, CV overestimates spiking irregularity in presence of rate fluctuations (Ponce-Alvarez et al., 2010); in contrast, as shown previously nΨ is robust to rate fluctuations (see Figure S2).

Specifically, we aimed to find some of the characteristics in the presynaptic network which control different sources of neural variability (i.e. nΨ and nRV). This is critical since the differences in the temporal dynamics of nΨ in vlPFC compared to other cortical areas or between subcortical and cortical areas, raise questions about underlying pre- and post-synaptic factors that derive spiking in different regions. Presynaptic network characteristics of the neural models are shown to have notable impact on the post-synaptic neuron variability (Cavallari et al., 2014; Deco & Hugues, 2012; Litwin-Kumar & Doiron, 2012). Rate correlation (Doiron et al., 2016; Salinas & Sejnowski, 2000) as well as spiking synchrony (Denève & Machens, 2016; Gómez-Laberge et al., 2016; Okun & Lampl, 2008; Salinas & Sejnowski, 2001) in presynaptic network are among these characteristics. Moreover, there are a wealth of studies suggesting a tight balance in cortical neurons (Denève & Machens, 2016; Renart et al., 2010) which might also affect the variability of the post-synaptic neurons. Here, we showed how statistical properties of balanced presynaptic network can reproduce some of the pattern observed in nΨ and nRV for a simple current based leaky integrate and fire (LIF) neuron.

We first show that the degree of trial-to-trial rate correlation between excitatory and inhibitory neurons (E and I Pools) in the presynaptic network can result in variations in nRV with minimal changes in nΨ in the post-synaptic neuron. Here we assumed two presynaptic pools, one excitatory and the other inhibitory sending efference to one LIF neuron (Figure 6a). The response variability of the LIF neuron was examined as a function of between pool trial-to-trial correlation in firing rate parameter (see methods). Figure 6b shows the firing raster of the LIF neuron for multiple scenarios from fully uncorrelated to fully correlated firing rates between E and I pools. As can be seen the increased rate correlation resulted in lower rate variability (nRV) in the post synaptic LIF neuron. Quantitative analysis showed that indeed nRV was high for zero correlation but decreased steadily for higher correlations. On the other hand, rate correlation had little effect on nΨ or nPPV (Figure 6d). The firing rate was also largely unaffected by the degree of E/I pool firing rate correlation (Figure 6c).

**Figure 6.**
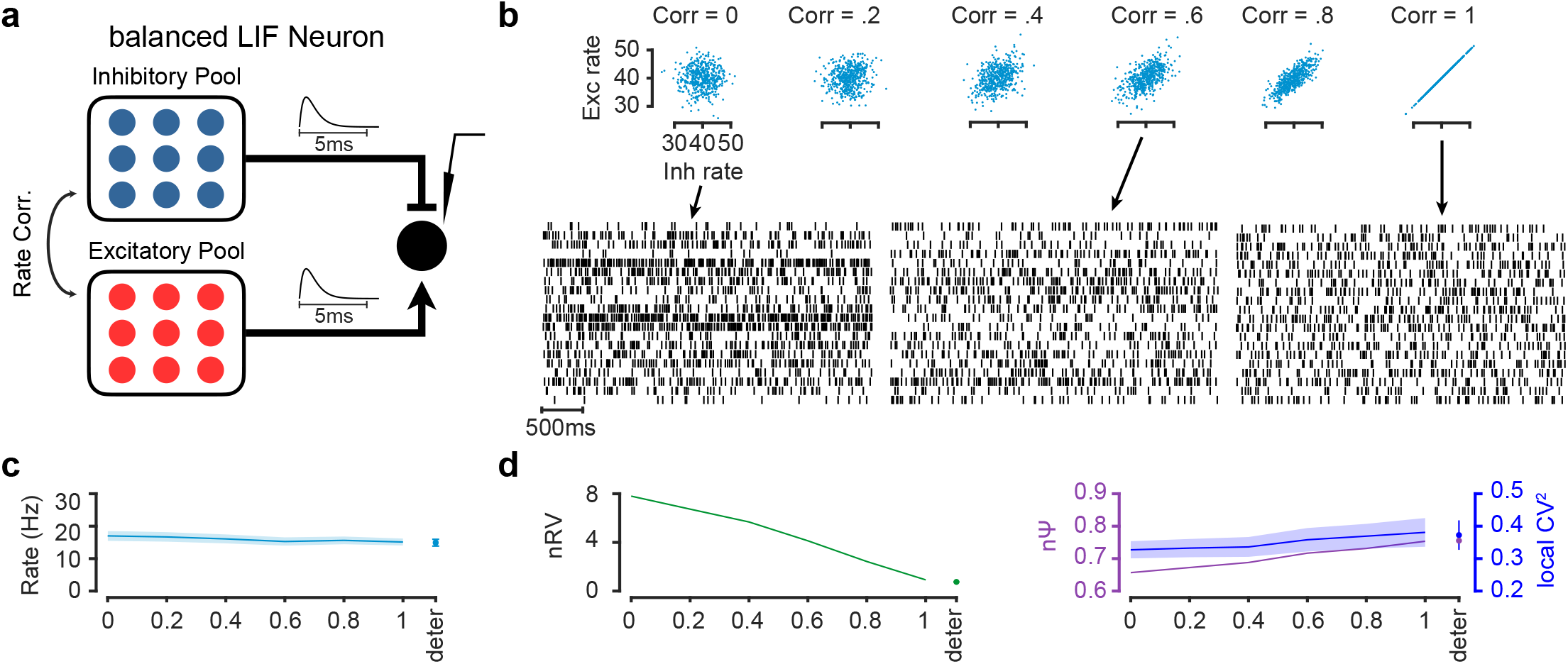
Firing rate correlation between balanced presynaptic excitatory and inhibitory pools controls post-synaptic nRV. (a) Schematic of the network model including 50 Excitatory (E pool) and 50 Inhibitory (I pool) Poisson neurons. The synaptic current is modeled using the same kernel for E and I connections. (b) E/I rate distributions of different trials (~ *N*(40,4)) with different correlation values (top) along with simulated sample rasters in three correlation conditions (correlations equal to 0, 0,6 and 1). (c) The effect of E/I rate correlation on the firing rate of post synaptic neuron. (d) The effect of E/I rate correlation on the nRV (left) and nΨ (right) of the post synaptic neuron. The effect on 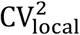 is shown alongside nΨ. Higher rate correlation leads to lower nRV in the post-synaptic neuron but had minimal effect on nΨ or 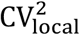. Firing rate, nRV nΨ and 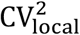 for nonrandom presynaptic firing rate (deter) is shown alongside for comparison in c-d. All error bars are SEM.

Next, it is shown that the degree of spiking correlation between and within E/I pools can result in reduction or enhancement of nΨ without affecting the nRV. Here we assumed two sub-pools (sub-pool #1 and #2) within each excitatory and inhibitory pools. In the first scenario, there was between pool spiking correlation (Figure 7a). In addition, a range of delays between spikes of excitatory and inhibitory neurons was considered (0, 2 and 4ms, aka tight balance (Denève & Machens, 2016)). Figure 7b shows simulated spiking realizations of E/I pools and the resulting spikes in the LIF neuron at 0.3 correlations (Smith & Kohn, 2008) and 0ms (left) or 4ms (right) delays. As can be seen, the spiking irregularity was higher at 0ms compared to 4ms delay in the post-synaptic neuron (Figure 7b top). Quantitative analysis confirmed the reduction in nΨ as a function of between pool spiking correlation for nonzero delays between E/I pools (Figure 7c). This manipulation did not affect nPPV nor the nRV. In this scenario the firing rate was also increased as the function of between pool spiking correlation. Similar results are found for nΨ, nRV, nPPV and rate by increasing the number of sub-pools (Figure S8).

**Figure 7.**
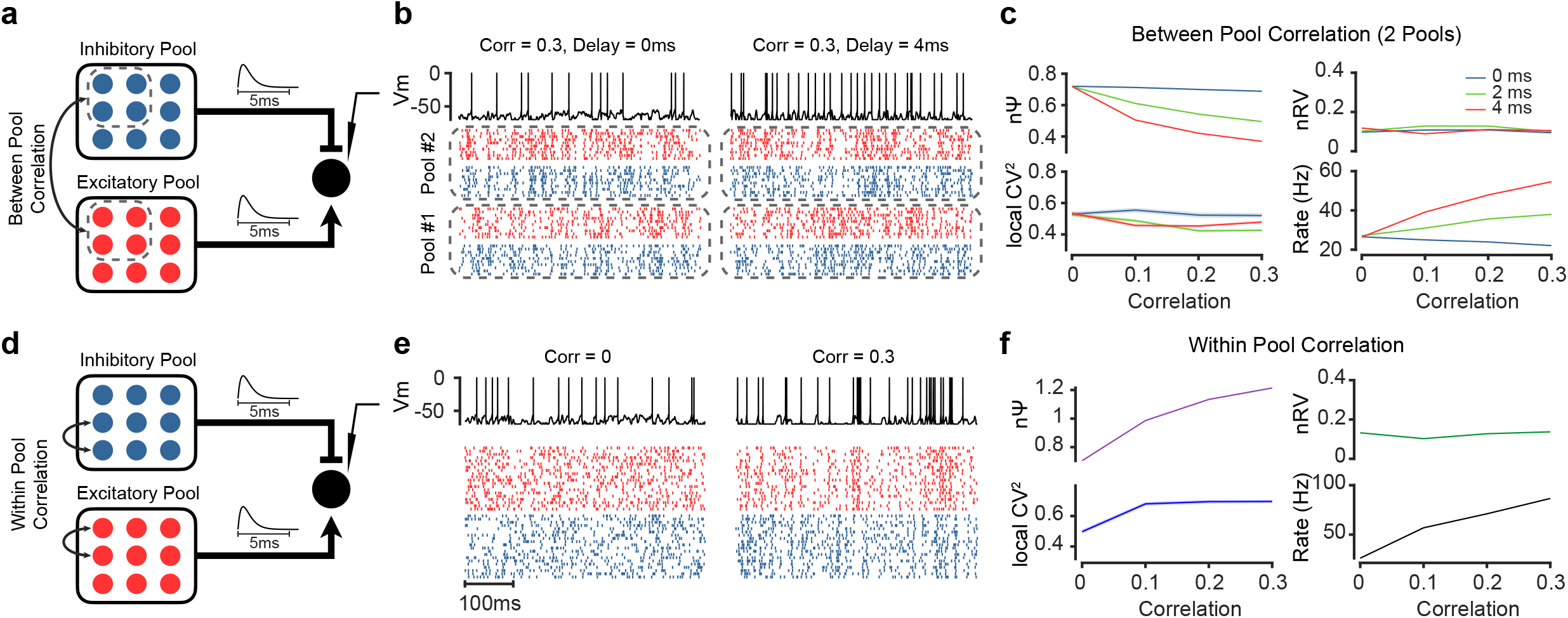
Spike correlation between and within balanced presynaptic excitatory and inhibitory pools controls post-synaptic nΨ. (a) Schematic of the network model consisting of 50 Excitatory (E pool) and 50 Inhibitory (I pool) Poisson neurons. Each pool is further divided to two sub-pools. Between pool spike correlation is imposed among sub-pools (between E and I sub-pool#1 and between E and I sub-pool#2). (b) Simulated spiking pattern in the E/I sub-pools 1 and 2 and the resulting spiking in the postsynaptic neuron when there is a 0.3 spiking correlation between E/I sub-pools with zero delay (left) and 4ms delay (right) between E/I neurons (see methods). (c) Stronger between pool spike correlation decreases nΨ for nonzero delays (2-4ms). nRV or 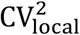 are not affected by between pool spike correlations. A concurrent increase in firing rate is also observed for stronger between pool spike correlation for nonzero delays (2-4ms). (d) The same network as (a) but with only within E and I pool spike correlation. (e) same format as b but for within pool correlations of zero (left) and 0.3 (right). (f) Stronger within pool spike correlation increases nΨ. nRV or 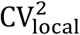 are not affected by within pool spike correlations. A concurrent increase in firing rate is also observed for stronger within pool spike correlation.

A different method to impose synchrony is to assume spike correlation within each E and I pools as well while excluding between pool correlation (Figure 7d). This scheme has been previously used to explain reduction of variability in response to partial inactivation of inputs from visual cortex (Gómez-Laberge et al., 2016). In this case increasing the within pool spiking correlation increased spiking irregularity as shown in the example simulated rasters (Figure 7e). Consistent with previous observations high correlations lead to bouts of excitatory activity, in absence of correlated inhibitory activity, which in part leads to burst like activity in the post-synaptic neuron (Gómez-Laberge et al., 2016). In contrast to between pool correlation, this increase in nΨ showed opposite polarity with firing rate. Once again, this within pool correlation also had no effect on nRV or nPPV.

Together these results show the plausibility of independent changes in nΨ and nRV as they were controlled by different presynaptic mechanism (rate correlation in Figure 6 vs spiking correlation in Figure 7). Notably in both cases nPPV as measured by 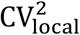 showed little changes similar to observations in the real data (Figure 2, 4, 5).

## Discussion

Neurons across the brain often show variable spiking pattern in response to the same stimulus or behavioral event (Carandini, 2004; M. M. Churchland et al., 2010; Shadlen & Newsome, 1994; Tomko & Crapper, 1974). This variability reflects both single cell level characteristics stemming from noisy sensory inputs (Schneeweis & Schnapf, 1999), stochastic synaptic transmission (Ribrault et al., 2011), plasticity (Feldman, 2009), and adaptation (Clifford et al., 2007); as well as network level properties including excitatory inhibitory balance (Turrigiano, 2011), attention and arousal levels (Fontanini & Katz, 2008), and neuromodulation effects (Marder, 2012). The total variability from these sources can be formulated into two additive components: (1) variability arising from random generation of spikes given ISI parameter which we referred to as spiking irregularity (nΨ), and (2) the trial-to-trial rate variation in ISI parameters which in the case of the renewal process stems from rate variability (nRV). By introducing new methods (FFA and Kass) for accurate estimation of nRV and nΨ in time, we were able to reveal concurrent changes in both components across multiple subcortical and cortical regions during the stimulus presentation in comparison to the spontaneous activity. Importantly, we showed that the previously reported quench in neural variability is solely due to reduction of nRV which was observed across all subcortical and cortical regions tested (A. K. Churchland et al., 2011; M. M. Churchland et al., 2010; Gómez-Laberge et al., 2016; Lombardo et al., 2018; Poland et al., 2019) (Figures 2, 4, 5). However, unlike what was previously assumed spiking irregularity was not constant but showed robust concurrent changes with nRV. nΨ, unlike nRV, did not necessarily decrease during stimulus presentation. In fact, IpN neurons in the cerebellum and SNr neurons showed robust increases in post-stimulus nΨ (Figures 2, 4). The increase in nΨ was also observable in the cortical neurons in the vlPFC region (Figure 5). The increase in spiking irregularity in SNr was so prominent that it canceled the reduction in rate variability resulting in the absence of usual quench in overall variability as measured by FF.

Specifically, for IPN neurons nΨ increased in response to the stimulus onset and also showed a sustained post-stimulus increase in FF after an initial dip (Figure 2). Purkinje cells (PC) which are presynaptic inputs to the IpN neurons are also shown to change their spiking regularity pattern in response to the stimulus onset. They also showed enhanced spiking irregularity due to inputs from molecular layer interneurons (Häusser & Clark, 1997; Payne et al., 2019; Shin et al., 2007). Although optogenetic manipulations in PCs have not shown sizable effects of nPPV on behavior (Payne et al., 2019), the changes in spiking regularity still has information about the presynaptic inputs and network level characteristics. The SNr neurons also showed increased nΨ in response to the stimulus onset invariant of the rate polarity such that post-stimulus FF remained almost constant despite the drop in nRV (Figure 4). The robust enhancement of nΨ in these subcortical regions compared to most cortical regions examined encourage further investigation on potential significance of spiking irregularity for the neural code and behavior and its underlying mechanism in different brain regions in future work.

Furthermore, nPPV, estimated by local measures of CV (Nawrot et al., 2008), has been previously used to explain the total variance caused by spiking irregularity (M. M. Churchland et al., 2010). However, our results showed that nPPV estimated by 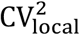 is not able to explain the total spiking irregularity observed in neural data (nΨ). We proposed several possible mechanisms that could explain this extra portion of spiking irregularity such as bursting, within trial rate noise as well as rate switching which were previously shown to be present in many brain regions and simulations based on neural networks (Deco & Hugues, 2012; Ermentrout et al., 2008; Ponce-Alvarez et al., 2010; Song et al., 2018). All these factors were shown to inflate nΨ beyond predictions of nPPV (Figure 2). In addition, network modelling results of between and within pool correlations in presynaptic E/I spikes were also shown to change spiking irregularity beyond inherent point process variability (Figure 7). This temporally correlated pre-synaptic activity has also been shown to be a way in which bombarding cortical neurons leads to post-synaptic spike irregularity simulations (Gómez-Laberge et al., 2016; Salinas & Sejnowski, 2000). Thus nΨ estimates relative to 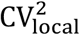 can be used to gauge the influence of sources of spike irregularity in addition to nPPV across brain regions. More research is required to both formulate the relation of these two spiking irregularity components as well as the underlying mechanisms responsible for the inflation of nΨ beyond nPPV and its possible role in neural processing and behavior.

We note that both methods used for estimation of variability components nRV and nΨ (i.e. FFA and Kass) require sufficient number spikes per time-bin (Figure S1). Thus, for neurons with low firing rates this would necessitates large time-bin and loss of temporal resolution in estimation of nRV and nΨ dynamics. Note that the requirement for the sufficient number of spikes per time-bin is due to the nature of doubly stochastic point process and exists even for idealized simulations with access to a large number of trial repetitions with the same parameter (i.e. empirical estimate, Figure S1). The relatively large time-bin used for estimation of nΨ may raise a concern about the changes in firing rate within time-bin to affect the variability estimates. However, our simulation results show that the spiking irregularity estimates are robust to such firing rate changes during the time-bin (Figure S2). As this figure illustrates, for different patterns of firing rate (Figure S2b-f) in comparison to the spontaneous activity (Figure S2a), the 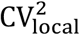 and nΨ remain largely unchanged. In case of large inhibition or excitation there could be some over and underestimation of our methods. In order to ensure the robustness of our main conclusions on the real neural data, we have implemented two additional strategies to control for variations in firing rate namely mean-matching and separately considering post-stimulus excitation, inhibition and null-responses for the variability analysis. Importantly, these additional controls resulted in virtually the same estimates of variability components and confirmed our main conclusions about relative roles of nRV and nΨ in total post-stimulus FF.

Different neurons show different spiking irregularity behaviors in vitro and in vivo; some of these characteristics are solely observed in vivo (network level variability) but not in vitro e.g. regularity of cortical pyramidal cells in vitro in spite of their irregular behavior in vivo. In contrast, in some other cases spiking irregularity is present in both in vivo and in vitro e.g. interneurons of the visual cortex (Englitz et al., 2008; Stiefel et al., 2013). In addition, blockage of tonic inhibitory inputs to the Purkinje cells of the cerebellum also leads to more regular spiking pattern (Häusser & Clark, 1997). In this work, we used LIF current based neural simulations to show effects of network input current properties on different sources of FF which models in vivo portion of variability. However, conductance based models are recently gaining ground as more viable alternatives to explain synaptic effects seen in real neurons (Cavallari et al., 2014; Richardson, 2004). The choice of synaptic model (current or conductance based) affects the spiking irregularity sensitivity of the post-synaptic neuron to the presynaptic firing rate. On the one hand, conductance-based neurons show persistent spiking irregularity roughly invariant of the presynaptic rate, consistent with our observations in V1, V2, MT, PMd cortical regions (Figure 5). On the other hand, the current-based neurons show sizable changes in their spiking irregularity with respect to the presynaptic firing rate which are consistent with our subcortical (Figure 2, 4) as well as vlPFC evaluations (Figure 5) (Cavallari et al., 2014). Thus, further evaluation of different sources of variability arising from these network models could provide new insights and diagnostic tools for proposing underlying neuronal and neural circuit mechanisms of the observed variability component across various brain regions.

Our results revealed a ubiquitous quench in post-stimulus nRV across many cortical and subcortical regions. Importantly we also find significant concurrent changes in nΨ which represents spiking irregularity in all areas examined. However, unlike nRV the pattern of post-stimulus changes in nΨ was heterogenous across regions. Previous work (M. M. Churchland et al., 2010) has indicated a robust reduction in network noise and a constant private noise during the stimulus. Our results do not address shared and private variability components across a population. However, the fact that both nΨ and nRV components showed concurrent post stimulus changes across neurons and had plausible network level implementations (Figure 6–7) suggest that they contribute to network level noise. On the other hand, nPPV seemed to be resistant to change in real data and thus may represent the private noise component for each neuron (Nawrot et al., 2008; Ponce-Alvarez et al., 2010). Interestingly, previous simulations and in vivo injections (Miura et al., 2006) have revealed that the balanced network of excitatory and inhibitory networks are plausible models explaining the constant nPPV phenomena as used in our simulations. Analysis of simultaneously recorded neurons using the Kass method (Vinci et al., 2016) which is readily available to address network level organization of nΨ and nRV across the population can further address factors underlying event related changes in neurons response variability components.

## Materials and Methods

### Subjects and datasets

#### Neural Data

MT and PMd data (n = 7031, 4264 sessions respectively) were recorded in macaques (M. M. Churchland et al., 2010) and are available online at https://churchland.zuckermaninstitute.columbia.edu/content/code.

The IpN data is from deep cerebellar nucleus of mice (ten Brinke, Heiney, Wang, Proietti-Onori, Boele, Bakermans, Medina, Gao, & De Zeeuw, 2017) and is available at (ten Brinke, Heiney, Wang, Proietti-Onori, Boele, Bakermans, Medina, Gao, & Zeeuw, 2017). We only used the first 270 sessions which were recorded during conditioning. V1 and V2 datasets (n = 1272, 296 respectively) are from anesthetized macaque monkeys in 5 different sessions (Zandvakili & Kohn, 2019). We used the session with largest number of neurons for each of these two regions and merged all different grating conditions. vlPFC and SNr data is recorded in macaques (Ghazizadeh et al., 2018; Ghazizadeh & Hikosaka, 2021). For all datasets, we removed low-rate neurons (sessions) with less than 3 spike/bin average for the given time-bin (number of used sessions are indicated in Figure 5).

#### Simulated spike trains

Spike trains in each trial were viewed as renewal point processes and ISIs were drawn from various distributions including Poisson, Gamma, and Inverse Gaussian processes (Figure 1a). For Figure 1b and Figure 3 Gamma distribution was used. In Figure 3d bursts were added to the spike trains by adding 5 consecutive spikes with ISI = 3ms on 5% of spikes. For rate noise (Figure 3e) we generated a subsampled white noise with Δ*t* = 50ms which was added to the rate of each trial randomly. The rate switching was simulated using a Markov Process with two fast and slow rate states (Figure 3f). The transition rates from slow to fast and fast to slow state are determined by *k_sf_* and *k_fs_* respectively (Song et al., 2018). The process on average is in the fast state with probability *p_f_*:

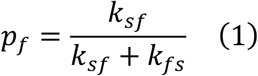

where we set *k_fs_* = 0.005ms^-1^ and *p_f_* = 0.4, hence the *k_sy_* ≈ 0.00 3ms^-1^. In order to constrain the average firing rate of the total process we set the slow rate based on the desired fast and average rate. The fast rate was set to *r_f_* = 25Hz and the slow rate would be determined using the following formulation:

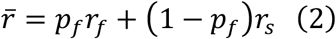

For average rate of 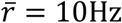 the slow rate equals *r_s_* ≈ 7.14Hz.

Between trial rate variability (RV) was imposed by drawing rates of different trials from a gaussian distribution with different standard deviations specified for each figure (Figure 1b, 3c).

#### Network simulations

We used LIF neuron with *N_E_* = *N_I_* = 50 neurons with the following formulation (Softky & Koch, 1993). We further set the parameters as in (Gómez-Laberge et al., 2016; Van Vreeswijk & Sompolinsky, 1996):

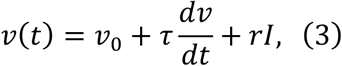

where *v_th_* = −55*mv, v*_0_ = −65*mv, v_min_* = −70*mv*, time constant *τ* = 20ms with synaptic resistance of *r* = 80*M*Ω and refractory period *t_ref_* = 1ms. The current *I* was:

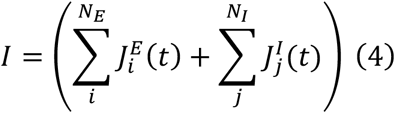

where 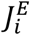 and 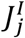 are presynaptic currents by each E/I pool neurons. We modeled the synaptic current response function by 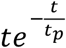, where *t_p_* = 0.5*ms* for both excitatory and inhibitory pools.

The presynaptic E/I pools (50 neurons/pool) were generated using correlated Poisson processes (Macke et al., 2009). We added a Laplacian distributed noise to the ISIs of the generated correlated neurons with mean zero and standard deviation of 2ms. For the correlation tightness we changed the mean to 2-4ms for inhibitory neurons in order to shift inhibitory neurons after excitatory ones on average. The rates of presynaptic pools in Figure 6 at each trial are generated using the bivariate normal distribution as illustrated in Figure 6b.

### Spike count variability decomposition

#### Fano Factor Asymptote (FFA) Method

Let *I*_1_, *I*_2_,…, *I_n_* denote the ISIs of a neural spike train. If we consider the distribution of each *I_j_* from an i.i.d. distribution the resulting point process is a renewal process. We leverage law of total variance and law of total expectation to decompose the spike count variability given the ISI distribution parameters in each trial *θ_i_*.

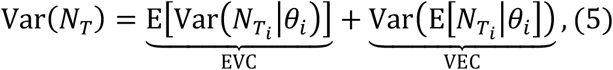

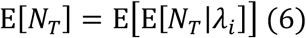

The first term in equation (5) represent the expected variance of count (EVC) while the second term represents variance of expected count (VEC) which was termed VarCE previously (A. K. Churchland et al., 2011).

We first derive equation (5) for doubly stochastic Poisson Process (exponentially distributed ISIs). For the ISIs from exponential distribution we have:

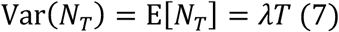

Substituting in (5) we have:

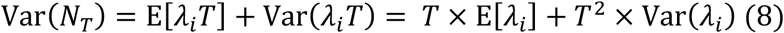

From (6) we have:

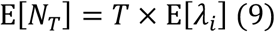

Moreover, from (7–9) we have:

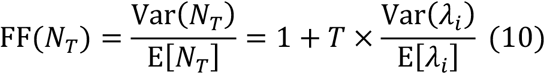

Hence, FF for the doubly stochastic Poisson process is a linear function of time bin *T*. As expected for stationary Poisson Process FF = 1.

In contrast to Poisson process, the aforementioned formulations do not have a closed form for general point processes. However, for renewal processes estimates of mean spike count and its variance can be calculated in the limit of large time bins and is shown to depend only on the mean and variance of the ISI distribution (Buckland & Cox, 1964). If 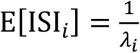 and 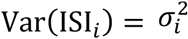, as such:

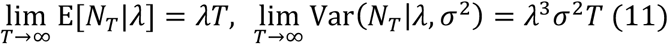

By substituting (11) in (5) we have:

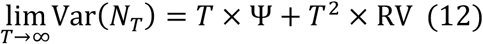

where

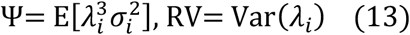

In addition, by substituting (11) in (6) the same equation (9) is derived for general renewal process. Fano factor (FF), the ratio of neural variability to its mean, is in effect a measure of neural count variability normalized by its mean. The decomposition presented in equations 12 allows one to parse FF into two components. Using equation (12) and (9), one can derive the asymptotic FF for a general renewal process in relatively large time bins:

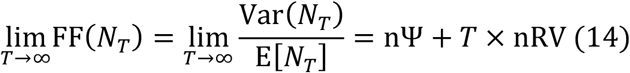

where

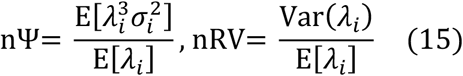

nΨ and nRV are normalized measures of Ψ and RV and also intercept and slope of FF in equation (14) respectively. Note that FF and E[*N_T_*] can be estimated empirically for specified time-bin (*T*). Note that the slope term in (10) and (14) representing the nRV are identical and invariant of the distribution type. Rather, they are proportional to changes of rate parameter across trials.

The FFA method works by first estimating the FF as a function of time-bin and then by fitting its linear asymptotic ([0.8:1] range of the estimation time-bin by default) behavior for which the slope represents nRV and the intercept measures nΨ (Figures 1a, c bottom, 2f, 4i, 5c).

Note that for a general doubly stochastic renewal process 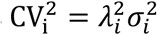, hence we have:

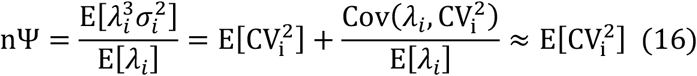

where the last approximation is by assuming that the CV^2^ as a measure of nPPV is not related to neuron’s firing rate; Note that while observations in cortical regions support this assumption (Ponce-Alvarez et al., 2010), it might not be true for all regions (Payne et al., 2019).

However, for the Gamma process in Figure 1b-e 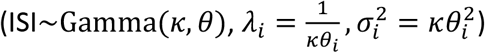, where *θ_i_* but not *κ* is randomly varying across trials, the equivalence of nΨ and CV^2^ is exact:

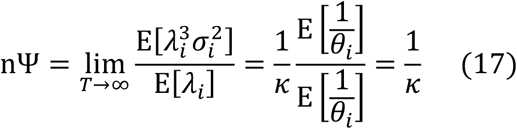

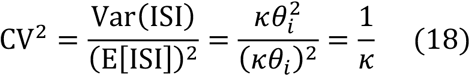

The same equality, with mild assumptions, is shown for the inverse-gaussian process in the last section (equations 25–27).

#### Modified Kass Method

The Kass method was originally introduced for decomposition of spike count correlation (SCC) between neurons into two components firing rate correlation (FRC) and within-trial point process noise (Vinci et al., 2016). This method was shown to be able to reliably extract the latent FRC from the observed SCC by a non-parametric approach. Obviously, the problem of spike count variance of a neuron can be considered as a special case of SCC between two neurons by substituting autocorrelations instead of cross correlations. Notably, the Kass method can be easily adapted to allow for a time-varying nΨ (aka *ϕ*), unlike previous methods which either took nΨ to be equal to one (Poisson process assumption) (Goris et al., 2014) or as a tuning variable to find relative RV respectively (A. K. Churchland et al., 2011).

Kass method requires a set of hyperparameters: (1) K which indicates the duration in which cross-correlation between two neurons is significant (duration of significant auto-correlation in our case) (2) bin-count (m) which indicates the number of bins that the time-bin is divided to (see (Vinci et al., 2016)). These parameters were set manually in the original implementation. In our implementation we used *m* = 100. For simulated data, the parameter K was set automatically for each neuron using its estimated shape parameter values (*κ*). We used equation (21) (see Local CV^2^ method) to find the shape parameter of the gamma process. Using the shape parameter and rate we first simulated a gamma process and empirically estimated the cross-correlation. K is estimated by the number of correlated lags (minimum correlation *α* ≥ 0.01). This method can be helpful in cases where there are low number of spikes per bin or limited number of trials. We further manually tuned the K parameter for real datasets to get more stable results.

#### VarCE Method

VarCE finds the largest constant *ϕ* such that VEC stays positive during the trial:

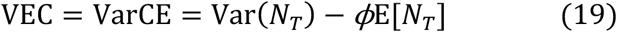

We used the code provided by (A. K. Churchland et al., 2011) to compute the VarCE and *ϕ*.

#### Local CV^2^ Method

Several local measures of CV, as a proxy of spiking irregularity, have been introduced in order to reduce sensitive to the rate variations in time (Davies et al., 2006; Holt et al., 1996; Miura et al., 2007; Ponce-Alvarez et al., 2010; Shinomoto et al., 2003). Theoretically for gamma process CV is only related to the shape parameter. Robust and unbiased estimation of the shape parameter could be achieved using the CV2 measure (Holt et al., 1996; Pachitariu et al., 2010). Normalized point-process variability (nPPV) which represents the inherent randomness of spike generation can be quantified using inverse of the shape parameter estimate which we refer to as 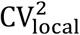. Although the gamma distribution of ISI has previously been examined by different evaluation on fitted distribution (Li et al., 2017; Maimon & Assad, 2009), even in non-renewal case, like occurrence of bursting, 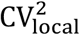 would persistently estimate the hidden gamma shape parameter (Pachitariu et al., 2010).

We used gamma shape estimator for non-stationary gamma processes to estimate the normalized point-process variability (nPPV) based on equation (17–18). CV2 is a measure of spiking irregularity which is robust to within trial non-stationarities (Holt et al., 1996). CV2(*i*) is computed by the following formulation:

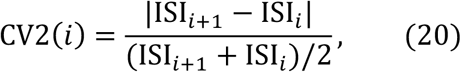

The shape parameter of the gamma process can be estimated using CV2 as follows (Pachitariu et al., 2010):

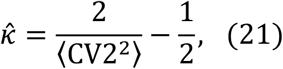

where, 〈.〉 denotes average. We then used this unbiased estimator to calculate the nPPV of gamma process which equals 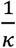 (equation 17–18) and which we refer to as 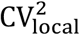:

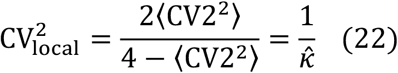

#### Mean Matching

Mean matching selects sub-population of neurons/conditions for each time-bin in trial period so as to the rate distribution remains constant for all time-bins. We used the code provided by the (M. M. Churchland et al., 2010) (Figure S5a–6).

#### Response polarity clustering

We used pre- and post-stimulus periods of the tasks to compute the average spike count for the two periods of each neuron. We then calculated the p-values for differences between pre- and post-stimulus rate of each neuron to cluster them into (1) excited (positive and p<0.05), (2) null (p>0.05), and (3) inhibited (negative and p<0.05) groups. We then used these clusters to calculate the average nΨ and nRV to test if the observed changes are due to rate fluctuations (Figure S5b-d, 7).

### Double stochasticity in the presented LIF network models

Approximation of network model in Figures 6 & 7 by simplifying assumption of integrate and fire (IF) model can be performed using Wiener process (no synaptic leakage). Hence, ISIs come from inverse gaussian distribution with following first and second statistical moments (for 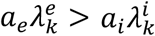).

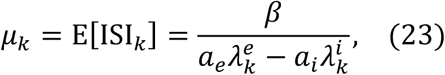

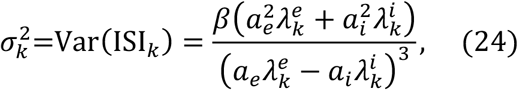

where, *β* represents threshold of post-synaptic IF neuron for spiking, 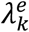 and 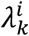 are excitatory and inhibitory firing rate of presynaptic neurons in trial *k*, and *α_e_* and *α_i_* are excitatory and inhibitory synaptic weights which are considered constant and equal to each other across trials (Tuckwell, 1988).

Hence, for large enough time bins, nΨ (or *ϕ*) can be computed by substituting (23) and (24) in (15) as follows:

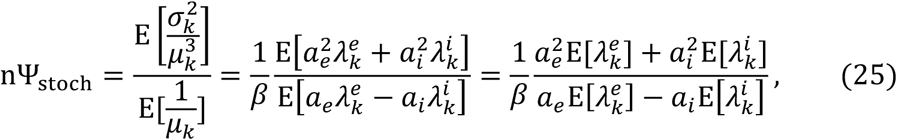

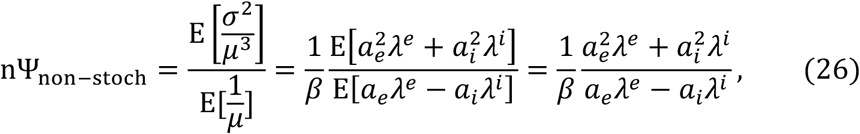

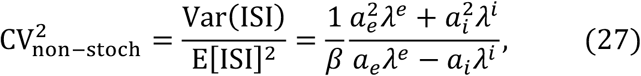

Equations (25–27) show that 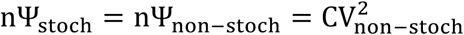. Given simplifying assumption of *a_e_* = *a_i_* = *a*, we have:

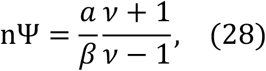

where, 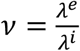 is the excitatory to inhibitory ratio.

## Acknowledgements

This research was supported by the Research Program at the Sharif University of Technology (Grant #G980703) and intramural grant at School of Cognitive Science, IPM.

## Author Contributions

S.F. devised the original ideas under A.G. supervision. S.F. and M.A.F did all the analysis and plots. All authors contributed to writing the manuscript.

## Competing Interests

The authors declare that they have no competing interests.

## Data and Code Availability

All data needed to evaluate the conclusions in the paper are present in the paper and/or the Supplementary Materials. All code and data are available upon request.

## List of supplementary materials

Figures S1–S8

## Supplemental Figure Captions

**Figure S1.**
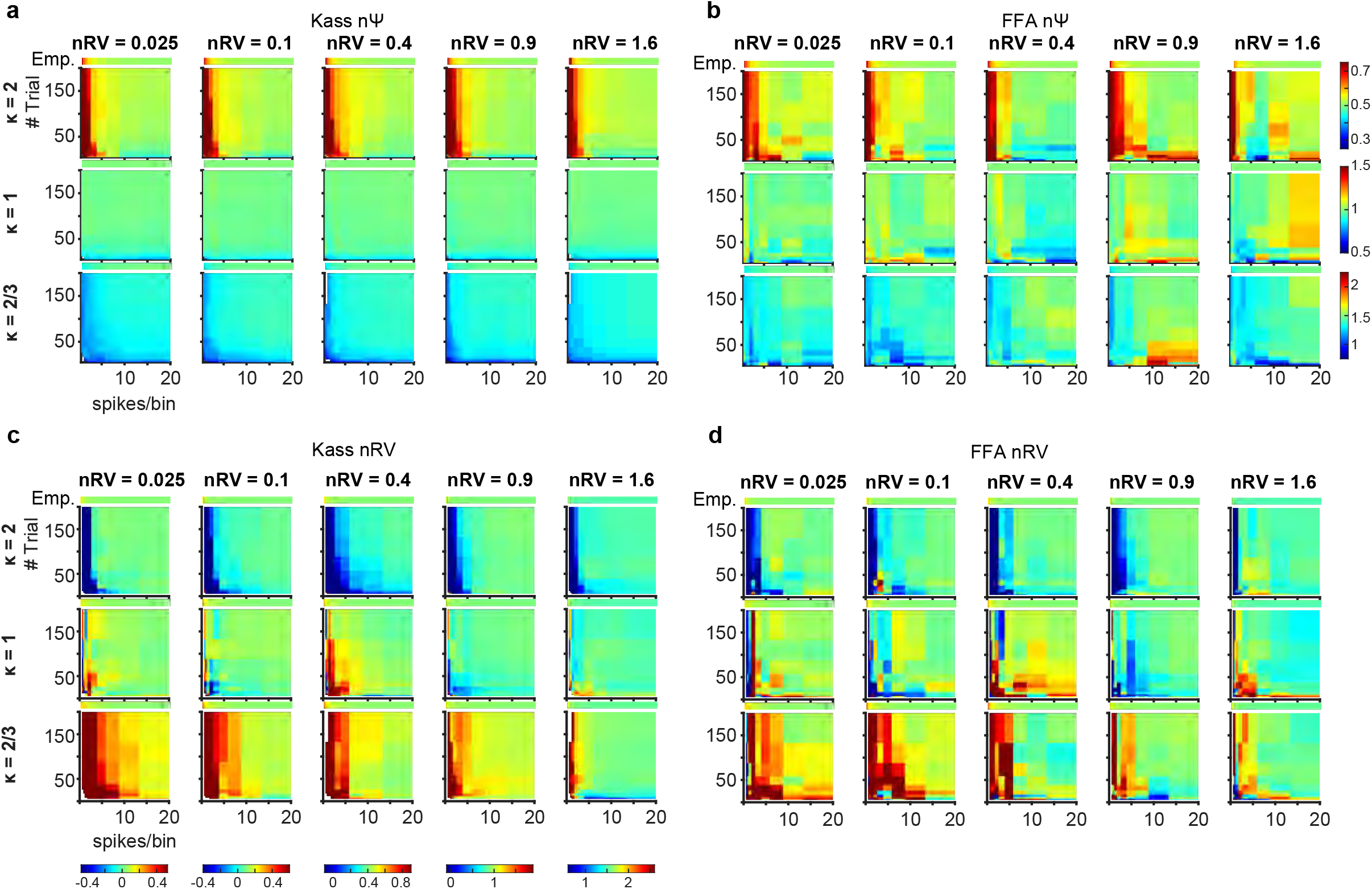
Dependence of nΨ and nRV estimates using Kass and FFA methods on the number of trials and the average number of spikes within the time-bin. (a) Each heat map shows estimated nΨ values for a gamma processe (10Hz) for different nRV (columns) and nΨ (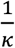, rows). X-axis shows average number of spikes within time-bin and y-axis shows number of trials. Color-code is adjusted such that green mean correct estimation, while red and blue mean over- and under-estimation of the true nΨ values. The heat bar over each square heatmap indicates the empirical estimates (best possible estimate given the repetition of each trial data). (b-d) same format as a but for estimated nRV values using the Kass method, for estimated nΨ values using FFA and for estimated nRV using FFA, respectively.

**Figure S2.**
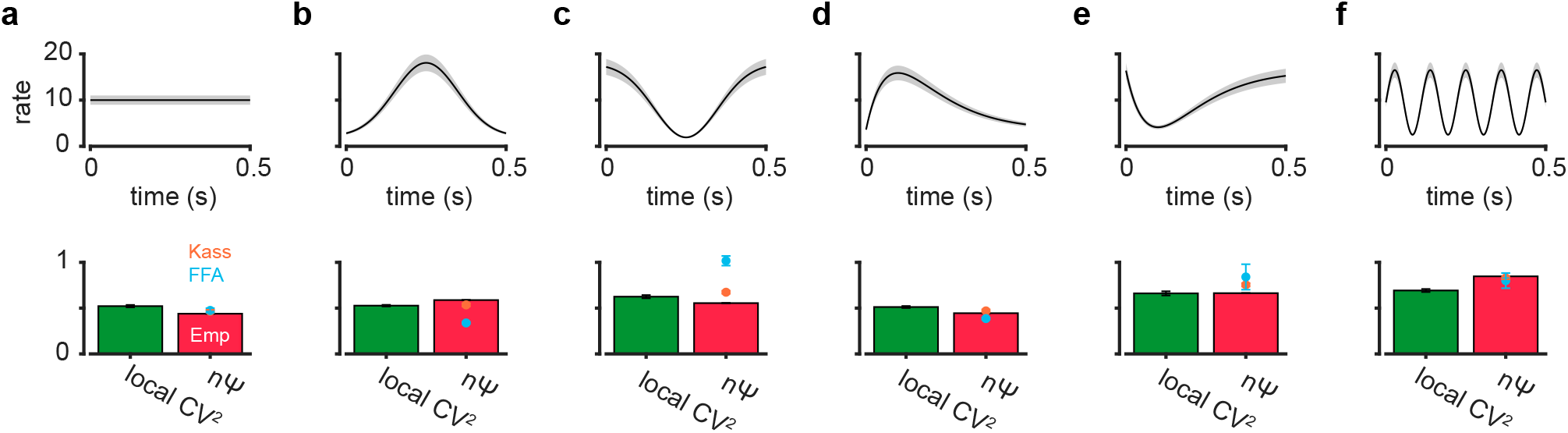
nΨ estimates are robust to changes in firing rate during the time-bin. (a-f, top) firing rate pattern within an time-bin of 500ms for a gamma process with *κ* = 2. The shading shows firing rate variation across trials which should only effect nRV not nΨ, as in Figure 3c. (a-f, bottom) Empirical estimates of nΨ (red bar) and the 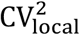 estimates (green bar) along with nΨ estimates by Kass and FFA method.

**Figure S3.**
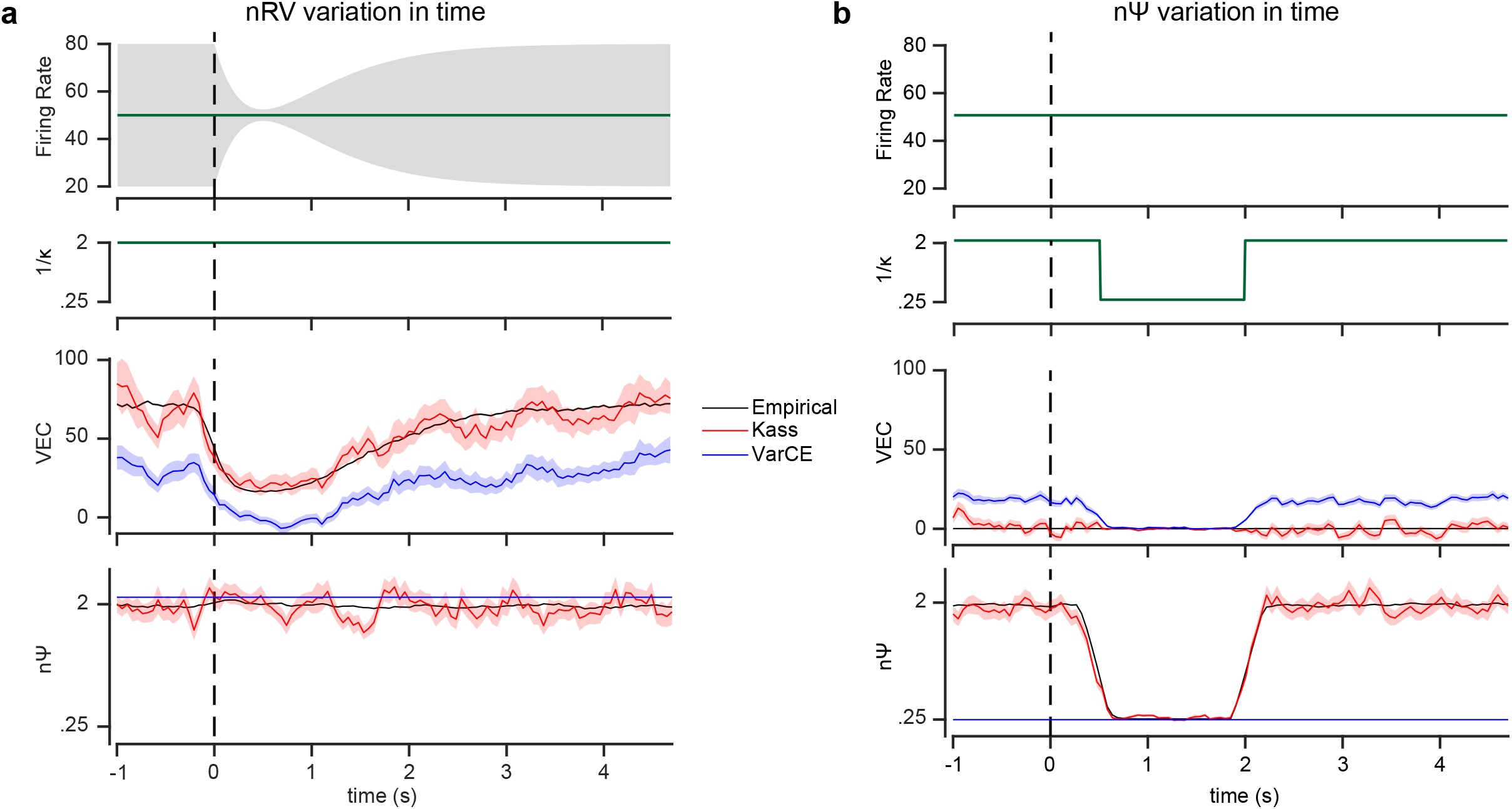
VarCE can fail when estimating VEC. (a) Similar to Figure 1b example with a time varying nRV component but a constant nΨ component (constant *κ*) (top two rows). Empirical estimates of VEC and nΨ (same as *ϕ*) along with estimates made by Kass and VarCE methods. (b) same format as (a) but for a case when there is no rate variation (VEC = *T*^2^ × RV = 0) but there is time varying spiking irregularity (changing *κ*). VarCE method assigns fluctuations caused by nΨ to VEC while Kass method correctly disentangles the two different sources.

**Figure S4.**
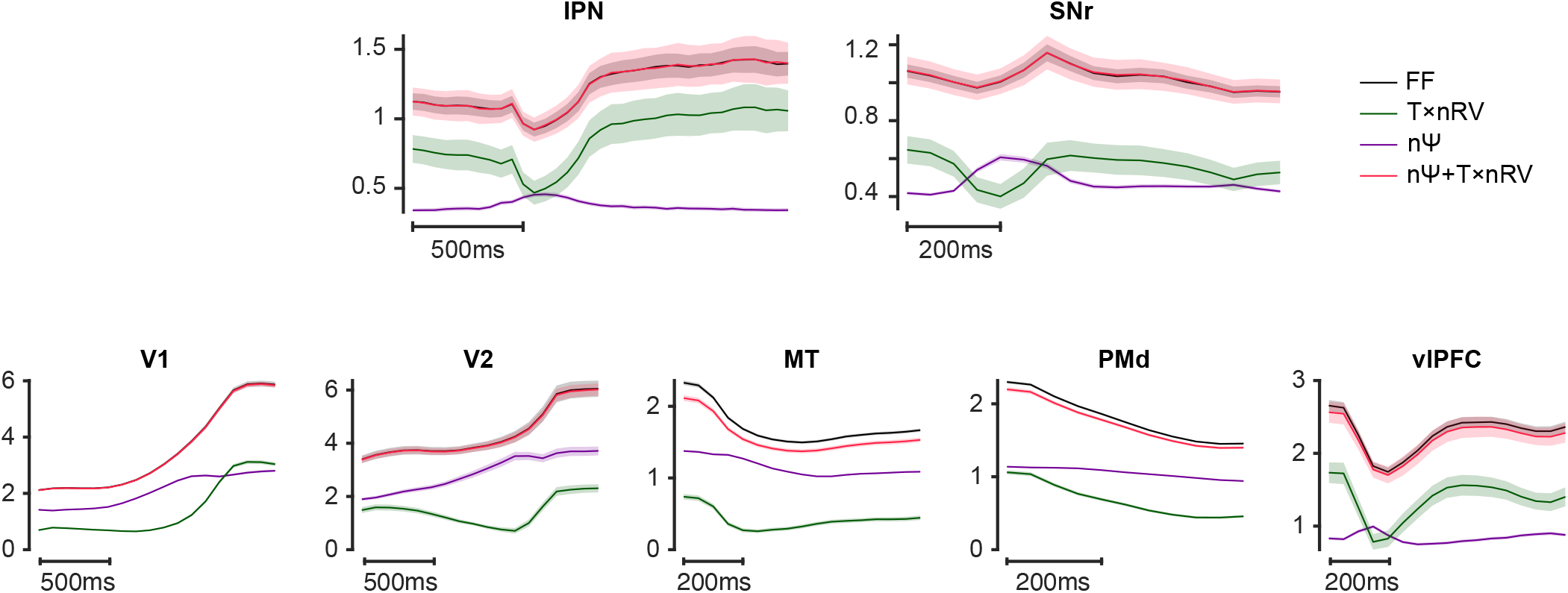
nΨ and nRV×*T* sum up to equal FF. Temporal dynamics of normalized spiking irregularity nΨ and normalized rate variability nRV estimates and the fact that they sum up to almost fully explain the FF dynamics. Results are shown for all subcortical and cortical regions analyzed in the main paper. The relative size and contribution of nΨ and nRV in driving the fluctuations in FF can also be examined.

**Figure S5.**
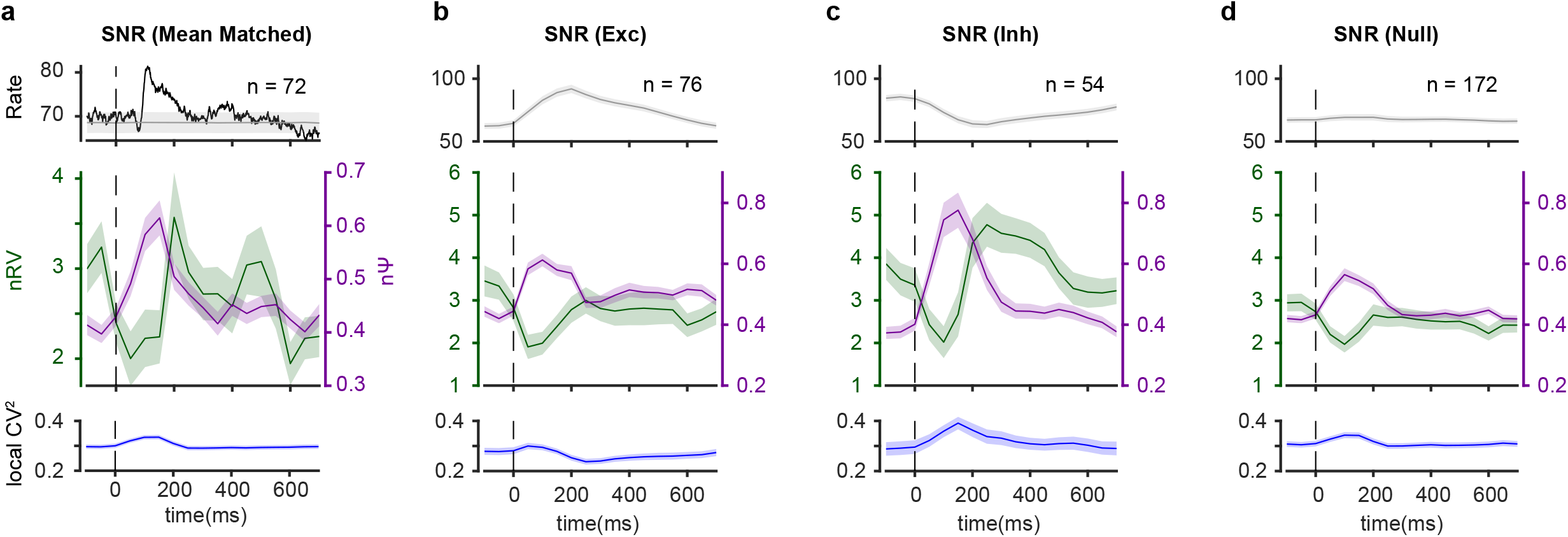
Robustness of nΨ and nRV pattern in SNr to variations in firing rate. (a) Mean-matching as well as clustering the SNr neurons to three groups with (b) excitatory response, (c) inhibitory response, (d) and null response yield similar patterns in nΨ and nRV estimates as shown in Figure 5. The gray curve in (a, top) plot shows the average and sem mean matched firing using a sub-selection of neurons for each time-bin. The gray curve in (b-d, top) is the PSTH evaluated with the same time-bin (200ms) as used for variability estimates nΨ and nRV.

**Figure S6.**
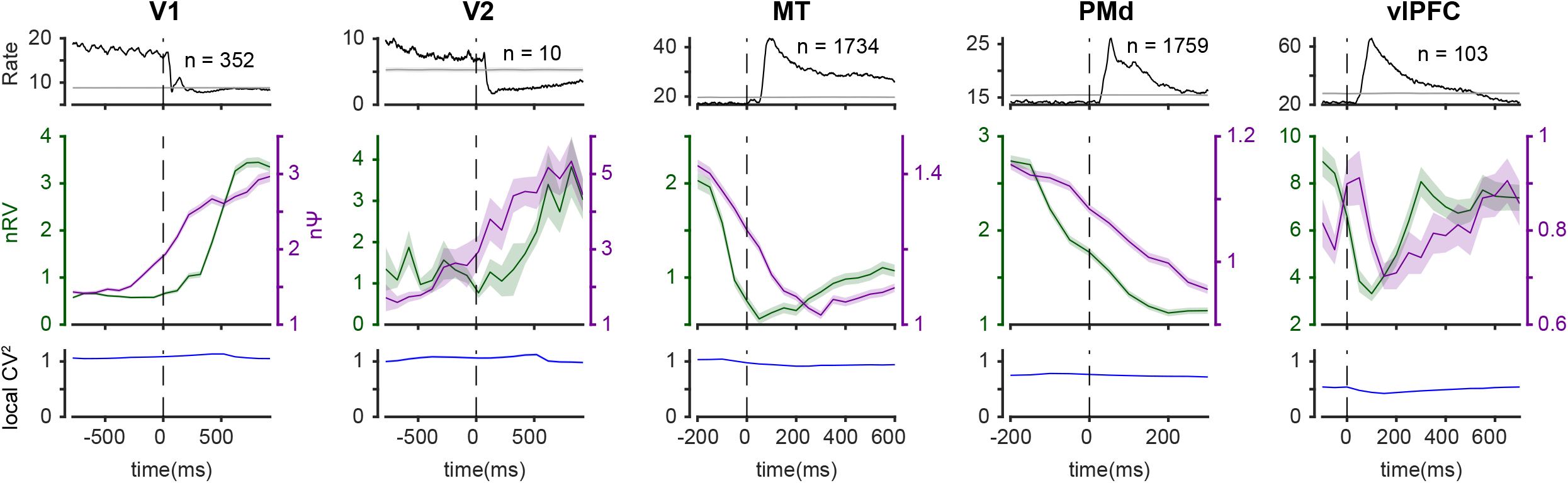
Robustness of nΨ and nRV pattern across cortical regions by mean-matching. Mean matching across cortical regions yield similar patterns in nΨ and nRV estimates as shown in Figure 4. The gray curved in the PSTH plot shows the average mean-matched firing rate.

**Figure S7.**
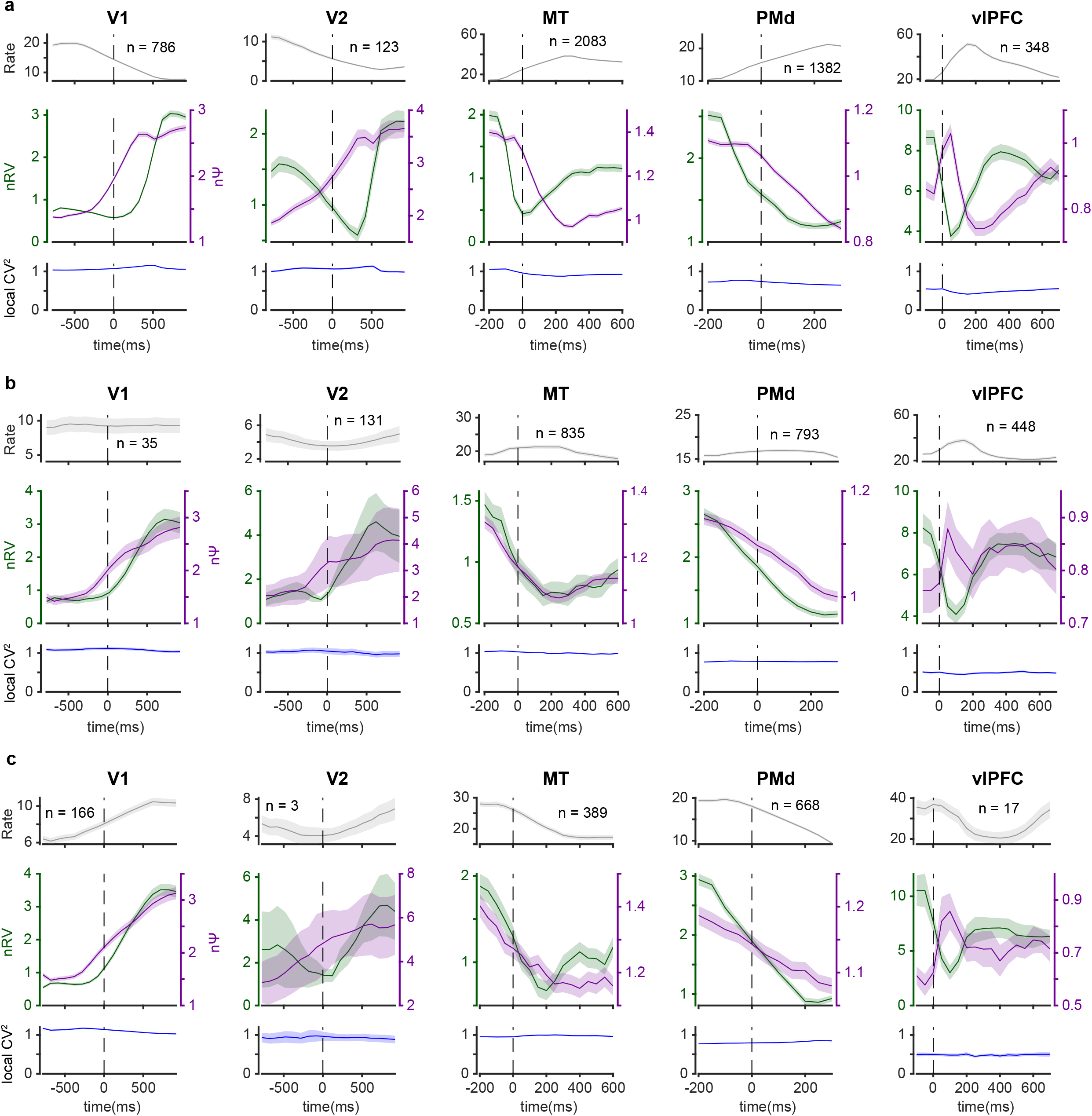
Robustness of nΨ and nRV pattern across cortical regions within response type clusters. Clustering cortical neurons to three groups with (a) excitatory response, (b) null response, (c) and inhibitory response yield similar patterns in nΨ and nRV estimates as shown in Figure 4.

**Figure S8.**
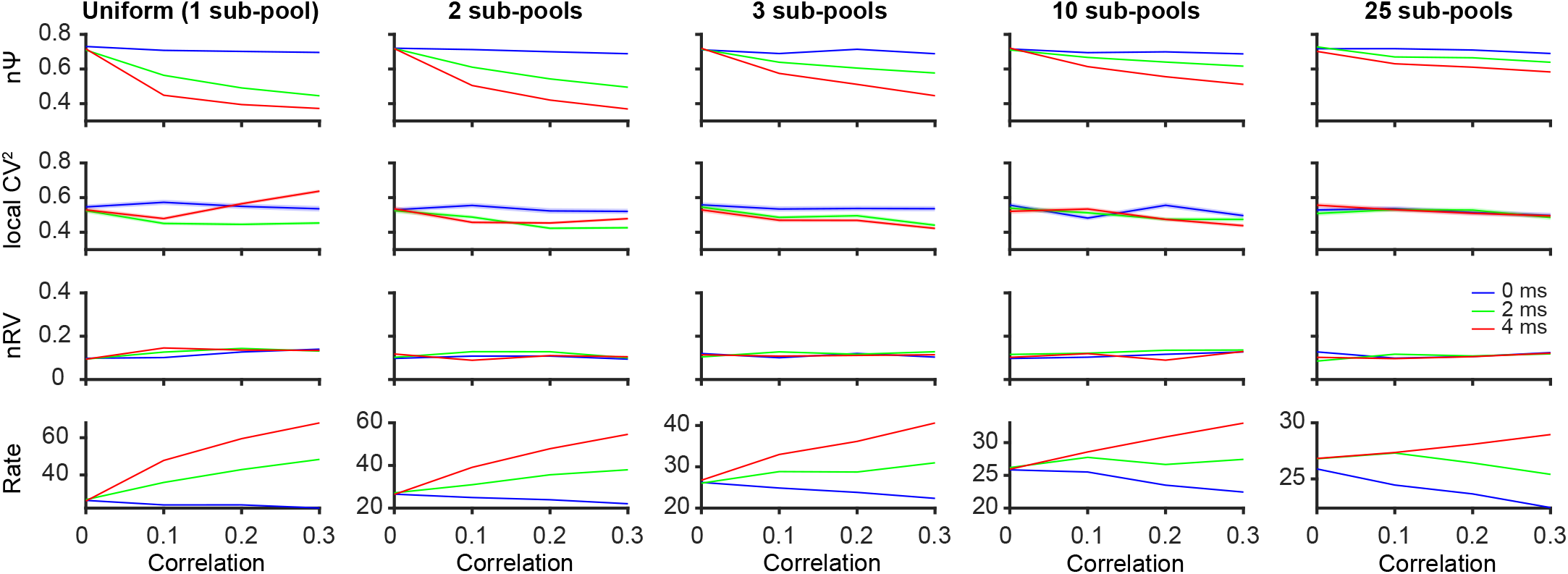
Effect of number of sub-pools on post-synaptic nΨ changes as a function of between pool correlation. nΨ and rate sensitivity to between pool correlation decreases as we increase the number of sub-pools (for constant number of neurons). nRV and 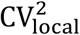 remain unchanged in all scenarios.

## Notes

### Competing Interest Statement

The authors have declared no competing interest.

